# Spen links RNA-mediated endogenous retrovirus silencing and X chromosome inactivation

**DOI:** 10.1101/2019.12.17.879445

**Authors:** Ava C. Carter, Jin Xu, Meagan Y. Nakamoto, Yuning Wei, Quanming Shi, James P. Broughton, Ryan C. Ransom, Ankit Salhotra, Surya D. Nagaraja, Rui Li, Diana R. Dou, Kathryn E. Yost, Seung Woo Cho, Anil Mistry, Michael T. Longaker, Robert T. Batey, Deborah S. Wuttke, Howard Y. Chang

## Abstract

Dosage compensation between the sexes has emerged independently multiple times during evolution, often harnessing long noncoding RNAs (lncRNAs) to alter gene expression on the sex chromosomes. In eutherian mammals, X chromosome inactivation (XCI) in females proceeds via the lncRNA *Xist*, which coats one of the two X chromosomes and recruits repressive proteins to epigenetically silence gene expression *in cis*^1,2^. How *Xist* evolved new functional RNA domains to recruit ancient, pleiotropic protein partners is of great interest. Here we show that Spen, an *Xist*-binding repressor protein essential for XCI^3-7^, binds to ancient retroviral RNA, performing a surveillance role to recruit chromatin silencing machinery to these parasitic loci. *Spen* inactivation leads to de-repression of a subset of endogenous retroviral (ERV) elements in embryonic stem cells, with gain of chromatin accessibility, active histone modifications, and *ERV* RNA transcription. Spen binds directly to *ERV* RNAs that show structural similarity to the A-repeat of *Xist*, a region critical for *Xist*-mediated gene silencing^8-9^. *ERV* RNA and *Xist* A-repeat bind the RRM3 domain of Spen in a competitive manner. Insertion of an ERV into an A-repeat deficient *Xist* rescues binding of *Xist* RNA to Spen and results in local gene silencing *in cis*. These results suggest that insertion of an ERV element into proto-*Xist* may have been a critical evolutionary event, which allowed *Xist* to coopt transposable element RNA-protein interactions to repurpose powerful antiviral chromatin silencing machinery for sex chromosome dosage compensation.

## INTRODUCTION

*Xist* is a 17kb long noncoding RNA that acts through specific interactions between its distinct RNA domains and nuclear effector proteins. The *Xist* RNA-associated protein complex was identified in 2015 using both genetic and affinity-based methods, and consists of multiple pleiotropic proteins, many of which are highly conserved throughout evolution and act on chromatin structure and gene regulation in myriad systems^3-7^. This suggests that *Xist* evolved the ability to bind these proteins in the eutherian mammals, coopting those which evolved initially to perform other epigenetic functions. *Xist* evolved in the eutherian clade through exaptation of a combination of coding genes that were pseudogenized, as well as transposable elements (TEs) that inserted into this locus. *Xist* contains six tandem repeat regions (A-F), all of which show sequence similarity to TEs, suggesting they arose from eutheria-specific TE insertions^9^. One of these is the A-repeat, which is essential for gene silencing. When this ∼500bp region is deleted, *Xist* RNA coats the X chromosome, but silencing and reorganization of the X does not follow^8,10^. The A-repeat sequence is thought to derive from the insertion and duplication of an endogenous retrovirus (ERV), a class of TEs present in many copies throughout the genome^9^. In general, lncRNAs are not well-conserved compared to protein-coding genes but are enriched for TE content, suggesting they may be able to rapidly evolve functional domains by exapting protein- and nucleic acid-binding activity from entire TEs that colonize their loci^11,12^. Understanding how the *Xist* RNA sequence was evolutionarily stitched together from these existing building blocks to gain protein-binding potential is of great interest towards understanding dosage compensation and lncRNA-mediated gene regulation genome-wide.

Spen (also known as SHARP, MINT) is a ∼400 kDa *Xist* RNA binding protein (RBP) that contains four canonical RNA binding domains, as well as a SPOC domain to facilitate protein-protein interactions. Spen is a co-repressor that binds to several chromatin remodeling complexes, including histone deacetylases (HDACs), and the NuRD complex^4,13^. Though now recognized for its central role in the eutherian-specific XCI process, Spen is an ancient protein that plays roles in gene repression during development in species including Drosophila and *Arabidopsis*, in addition to mice and humans^13-17^. Spen binds directly to the A-repeat of *Xist* RNA and *Spen* inactivation abrogates silencing of multiple X-linked genes, suggesting that the RNA-protein interaction between the A-repeat and Spen is an early and essential step in XCI^3,4,6,18^.

## RESULTS

To test the effect of Spen loss on gene regulation and chromosome accessibility during XCI and genome-wide during development, we performed ATAC-seq^19^ in haploid mouse embryonic stem cells (mESCs) harboring a doxycycline-inducible *Xist* transgene either in the wild-type (WT) context or with a full deletion of *Spen* (*Spen* KO)^6^ (**Figure 1a**). Following 48 hours of *Xist* induction, WT cells demonstrated loss of chromatin accessibility at the majority of loci on the X chromosome (**Figure 1b-c**; Extended Data Figure 1a,b). In two independent *Spen* KO mESC clones, we found no X chromosome site that is reproducibly silenced upon *Xist* induction, suggesting that Spen is absolutely required for gene silencing at the level of chromatin accessibility (**Extended Data Figure 1b**). This complete failure of XCI, combined with Spen’s direct binding to the A-repeat region^3,4,6^, confirms that Spen’s recruitment to the inactivating X chromosome is early and essential for XCI.

**Figure 1:**
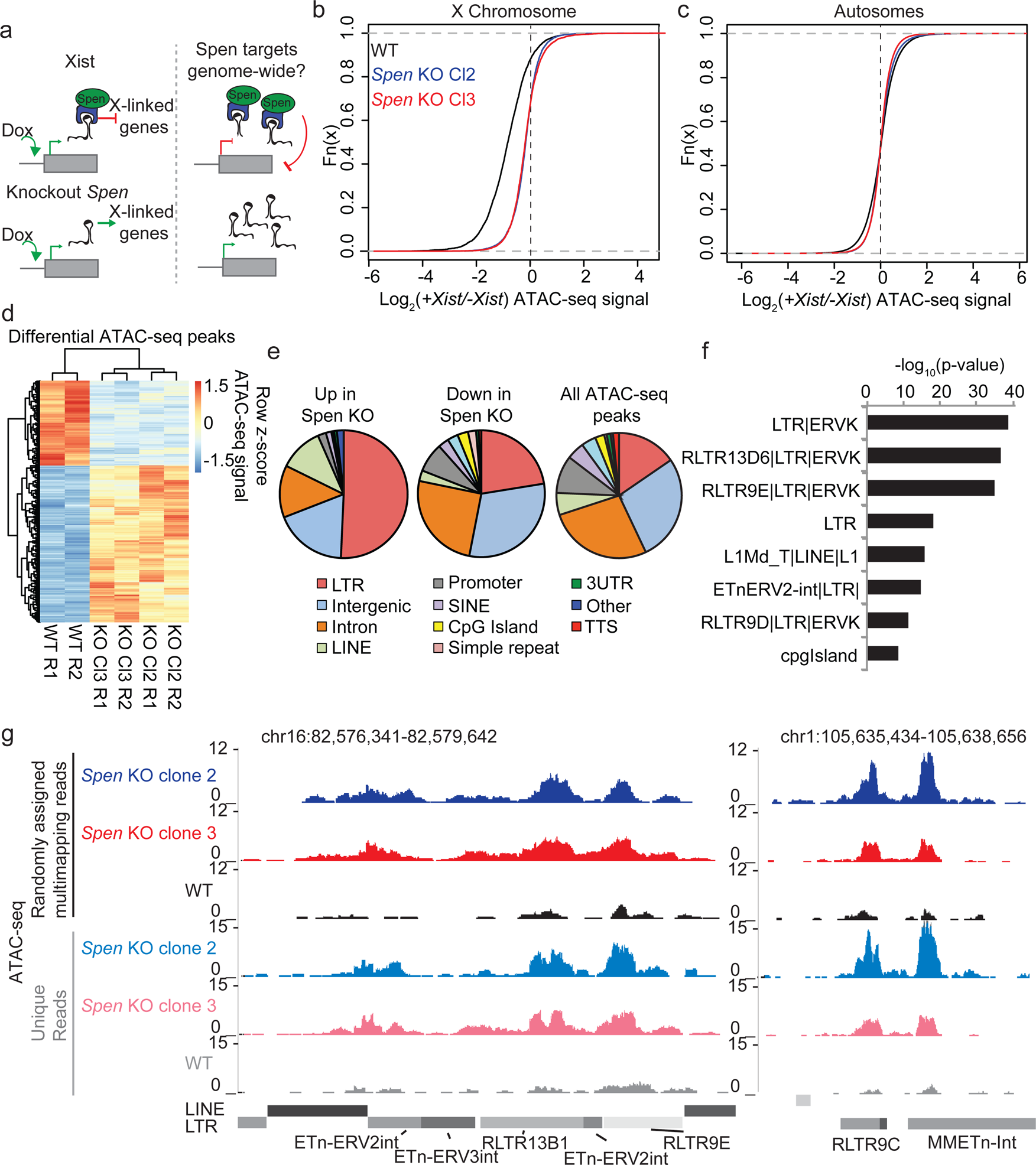
*Spen* knockout blocks XCI chromosome-wide and leads to derepression of autosomal ERVs. a. Diagram of experimental set up. *Spen* KO mESCs are cultured +/-Doxycycline for 48 hours to induce *Xist* expression and then ATAC-seq is performed. We ask what sites are de-repressed in *Spen* KO compared to WT cells both during XCI and on the autosomes. b. Log_2_ ratio of ATAC-seq signal in +Dox and –Dox samples at all peaks on the X chromosome in WT and *Spen* KO mESCs. Two independent *Spen* KO mESC clones are shown. c. Same as in b for all autosomal peaks. d. Heatmap showing ATAC-seq signal for all differential peaks between WT and *Spen* KO mESCs. Values depict the z-scored value for normalized ATAC-seq reads within each peak for WT, *Spen* KO Cl2, and *Spen* KO Cl3 cells (with 2 replicates each). e. Pie charts depicting the proportion of ATAC-seq peaks that cover each genomic region annotation. Annotations are derived from HOMER. f. Genome Ontology enrichment (by HOMER) for all sites gaining accessibility in *Spen* KO mESCs compared to WT (*n*=288). Plotted is the –log_10_ p-value for enrichment. *P* values (one-sided binomial) are accounted for multiple hypothesis testing using the Benjamini-Hochberg correction. g. Two example ERVK regions on chromosome 16 (left) and chromosome 1 (right) showing a gain in chromatin accessibility specifically in two clones of *Spen* KO mESCs. Shown are tracks of uniquely mapping reads (top) and tracks where multimapping reads are randomly assigned to one location (bottom).

Despite recent focus on Spen’s role in XCI, Spen is an ancient protein that is known to act as an important RNA-protein scaffold in developmental processes in many species^13-17^. Thus, how *Xist* evolved the ability to recruit Spen is of great interest. We hypothesized that understanding Spen’s role in autosomal gene regulation and its target specificity might lend clues into how Spen was exapted to participate in the Eutherian-specific process of XCI (**Figure 1a**). To assess the effects of Spen loss on autosomal gene regulation, we compared ATAC-seq data genome-wide in WT and *Spen* KO mESCs. We found that there are 288 sites on the autosomes that gain accessibility in two clones of *Spen* KO mESCs, compared to only 147 sites that lose accessibility (**Figure 1d**). This observation is consistent with chromatin de-repression in the absence of Spen’s repressive function. The DNA elements that were more accessible in *Spen* KO mESCs are almost all distal to transcriptional start sites (TSS), in contrast to unchanging peaks and those that are less accessible, which encompassed both promoters and distal sites (**Extended Data Figure 1c**). This indicated that DNA elements regulated by Spen on the autosomes are not found at gene promoters, but rather are found in heterochromatic, gene-poor regions.

To better understand Spen’s autosomal targets, we performed genome ontology enrichment for the sites that are de-repressed in *Spen* KO mESCs using HOMER. While the set of sites confidently gaining accessibility in both *Spen* KO clones is relatively small, it showed a striking enrichment for TE-derived long terminal repeat (LTR) elements (**Figure 1e**; **Extended Data Figure 1d**). When we looked more closely at subsets of these annotations, we found an enrichment specifically for endogenous retrovirus K (ERVK) TEs in these sites, which is reproduced in two independent *Spen* KO clones (**Figure 1f**). These ERVKs are enriched specifically for LTR elements in the RLTR13, RLTR9, and early transposon (ETn) families (**Figure 1f**). ERV-derived sequences in the genome have recently been shown to play important roles in genome regulation in embryonic cell types, serving as binding sites for transcription factors, indicating that Spen loss may activate ectopic regulatory regions^21-23^. Because sequencing reads coming from TEs in the genome do not map uniquely, it is difficult to accurately quantify reads coming from a given ERV subfamily while also mapping them to the correct element. We thus mapped ATAC-seq reads to the genome, while either keeping only uniquely mapping reads, or allowing multi-mapping reads to randomly map to one location. Both of these methods revealed the same trend, demonstrating the de-repression of a subset of ERVK elements in *Spen* KO mESCs (**Figure 1g**).

The observation that TE-derived elements were activated in *Spen* KO mESCs was intriguing, given that the A-repeat region of Xist is itself believed to be derived from an ancient TE insertion^9^. It has been posited that TEs, upon insertion into pseudogenes or non-coding loci maycontribute functional protein-binding domains to non-coding RNAs^11^. Indeed, noncoding RNAs, including Xist, are enriched for TE content^12^. Furthermore, it is known that in a very distantly related species, the model plant *Arabidopsis*, the Spen homologs *fca* and *fpa* bind to and regulate transcription of TEs in the genome^17^. Therefore, Spen’s ability to regulate ERVK, may explain Spen’s ability to recognize the A-repeat and interact directly with it.

In mESCs, the majority of TEs, including ERVKs, are silenced by histone H3 lysine 9 trimethylation (H3K9me3). When members of the TE silencing machinery, such as Kap1 and Setdb1, are inactivated, H3K9me3 is lost, and these TE insertions can be expressed and function as ectopic promoters and enhancers^25-27^. We hypothesized that in the absence of Spen and its protein partners, these ERVKs would lose H3K9me3 and gain histone modifications associated with active gene expression^2,4,13^.

Indeed, ChIP-seq experiments revealed that DNA elements gaining accessibility in *Spen* KO showed a dramatic loss of H3K9me3 and a gain of both H3K27Ac and H3K4me3 marks (**Figure 2a,b**). Conversely, H3K9me3 peaks that are lost in *Spen* KO mESCs are enriched for ERVKs, and sites that gain the enhancer mark H3K27Ac are enriched for ERVKs including ETn elements (**Figure 2c,d**). This demonstrates that at the level of chromatin modifications, a subset of ERVK elements are de-repressed when Spen is lost in mESCs. To test whether these ERVK elements are upregulated at the level of transcription, we performed RNA-seq in WT and *Spen* KO mESCs (**Extended Data Figure 2a-c**). At the RNA level, the ERVK elements that gain accessibility also have higher expression in *Spen* KO mESCs (**Figure Figure 2b,e**). Genome-wide there is a very small increase in expression of all ERVKs, particularly the class of ETnERV2s, indicating that only a subset of these families are affected by the loss of Spen in mESCs (**Figure 2f,g**).

**Figure 2:**
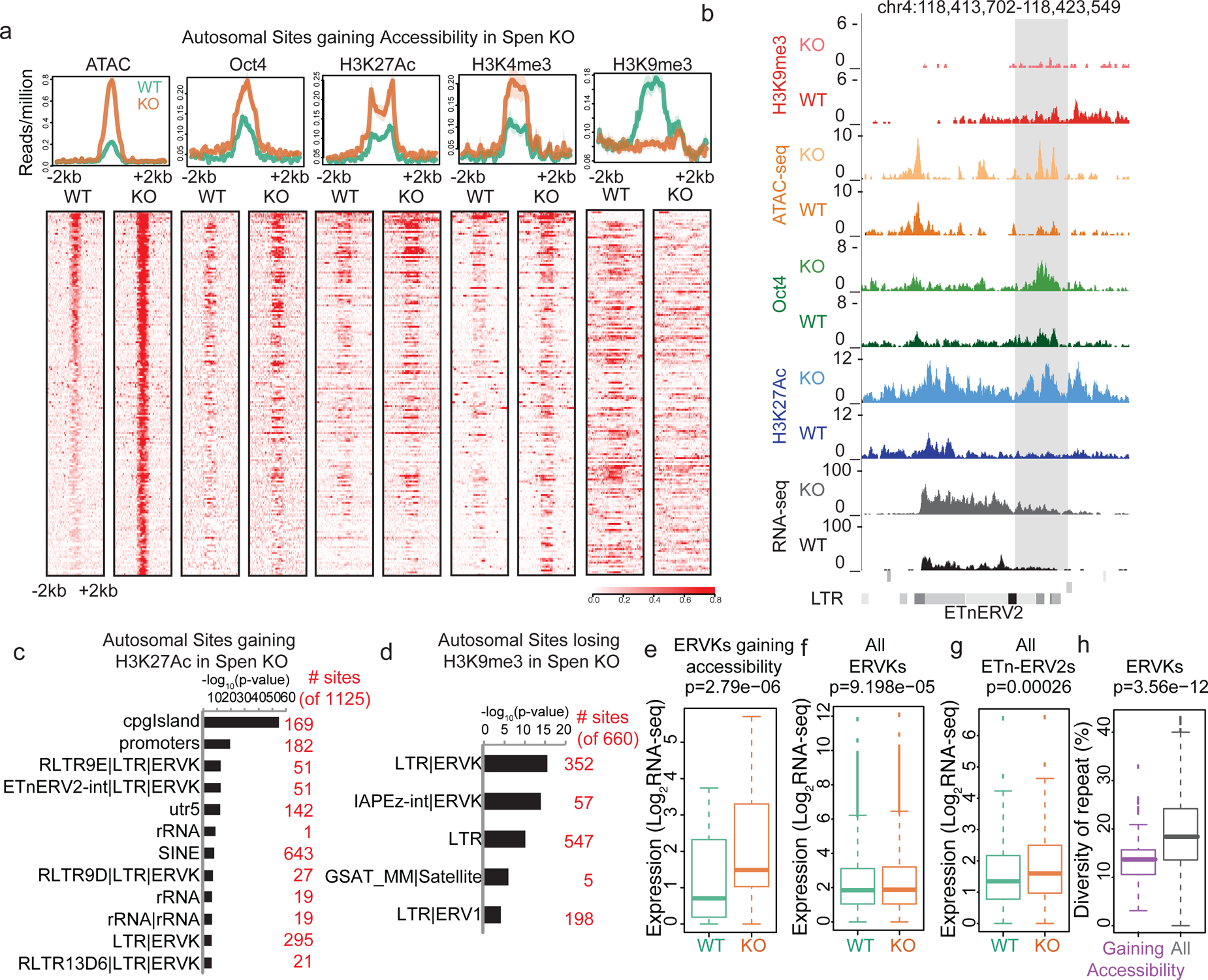
*Spen* knockout leads to gain of histone acetylation and loss of H3K9me3-mediated repression at ERVKs. a. Average diagrams and heatmaps showing the read count per million mapped reads for ATAC-seq and ChIP-seq plotted over all ATAC-seq peaks that gain accessibility in *Spen* KO mESCs (*n*=288) (+/-2kb). From left to right are ATAC-seq, Oct4 ChIP-seq, H2K27Ac ChIP-seq, H3K4me3 ChIP-seq, and H3K9me3 ChIP-seq. Data is shown for WT mESCs (green) and *Spen* KO clone 2 mESCs (orange). For the heatmaps, each row represents one peak. b. An example of an ETnERV2 element on chromosome 4 that shows a gain of chromatin accessibility, RNA expression, H3K27Ac ChIP-seq, Oct4 ChIP-seq, and a loss of H3K9me3 ChIP-seq in *Spen* KO mESCs compared to WT. c. Genome Ontology enrichment (by HOMER) for all sites gaining H3K27Ac ChIP-seq signal in *Spen* KO mESCs compared to WT (*n=*1125 sites). Plotted is the–log_10_ p-value for enrichment. *P* values (one-sided binomial) are corrected using the Benjamini-Hochberg correction. d. Genome Ontology enrichment (by HOMER) for all sites losing H3K9me3 ChIP-seq signal in *Spen* KO mESCs compared to WT (*n=*660 sites). Plotted is the –log_10_ p-value for enrichment. *P* values (one-sided binomial) are corrected using the Benjamini-Hochberg correction. e. Expression of ERVKs gaining accessibility in *Spen* KO mESCs in WT mESCs (green) and *Spen* KO mESCs (orange). Shown is the log_2_ transformed normalized RNA-seq read counts. For all boxplots, the thick line represents the median, while the box gives the IQR. *P* values are calculated using a two-tailed T test. f. Same as in d for all ERVKs, genome wide. g. Same as in e for all ETn-ERV2s. h. Diversity of repeat for all ERVK elements that gain accessibility in *Spen* KO mESCs (purple) and all ERVK elements (gray). Percent diversity represents the percentage of the sequence that has diverged from the ancestral TE insertion. Percentages come from RepeatMasker.

Retrotransposons are parasitic elements whose insertion and propagation in cells deplete cellular resources and disrupt endogenous gene expression. In order for the cell to handle these parasitic elements, pathways have evolved to shut down these transposons at the level of the DNA locus and the RNA transcript. ERVK insertions that are more similar to the original TE sequence pose more of a threat to cells, as they are more likely to be mobile and able to replicate. Thus, we compared the sequence diversity of the ERVKs upregulated in S*pen* KO to those not upregulated in *Spen* KO, and found that those repressed by Spen have diverged less than those that are not repressed by Spen (**Figure 2h**). The lower diversity indicates that Spen targets more recent and active ERVK insertions. Furthermore, we found that genes involved in the innate cellular immunity pathway, which responds to retroviral RNA presence, are upregulated in *Spen* KO mESCs (**Extended Data Figure 2d**). These data suggest that Spen represses young, more intact ERVKs, and when Spen is inactivated, the cell responds to the presence of those RNA transcripts to attempt to silence them.

TE insertions are enriched for transcription factor binding motifs, and it has been suggested that insertion of many elements of the same family within the genome may have led to the evolution of transcription factor-based regulatory networks^21^. ERVK, and more specifically ETn, elements are enriched for binding sites for Oct4, one of the most critical transcription factors for regulation of the pluripotent state^21,28^. Oct4 plays a critical role in balancing the expression of pro-self renewal and pro-differentiation genes in mESCs. We find that the Oct4 binding motif is enriched at DNA elements gaining accessibility in *Spen* KO mESCs, and Oct4 occupancy increases at these loci as shown by ChIP-seq (**Figure 2a**). The ability of this core stemness factor to bind to these TE elements suggests that their misregulation in *Spen* KO may affect the self-renewal or differentiation programs within these mESCs. GO term analysis revealed that genes downregulated in *Spen* KO mESCs are enriched for early developmental terms, indicating that these cells may be deficient in early lineage commitment (**Extended Data Figure 2f**). We found that *Spen* KO mESCs downregulated specific genes associated with early differentiation and upregulated genes associated with self-renewal (**Extended Data Figure 3a**).

**Figure 3:**
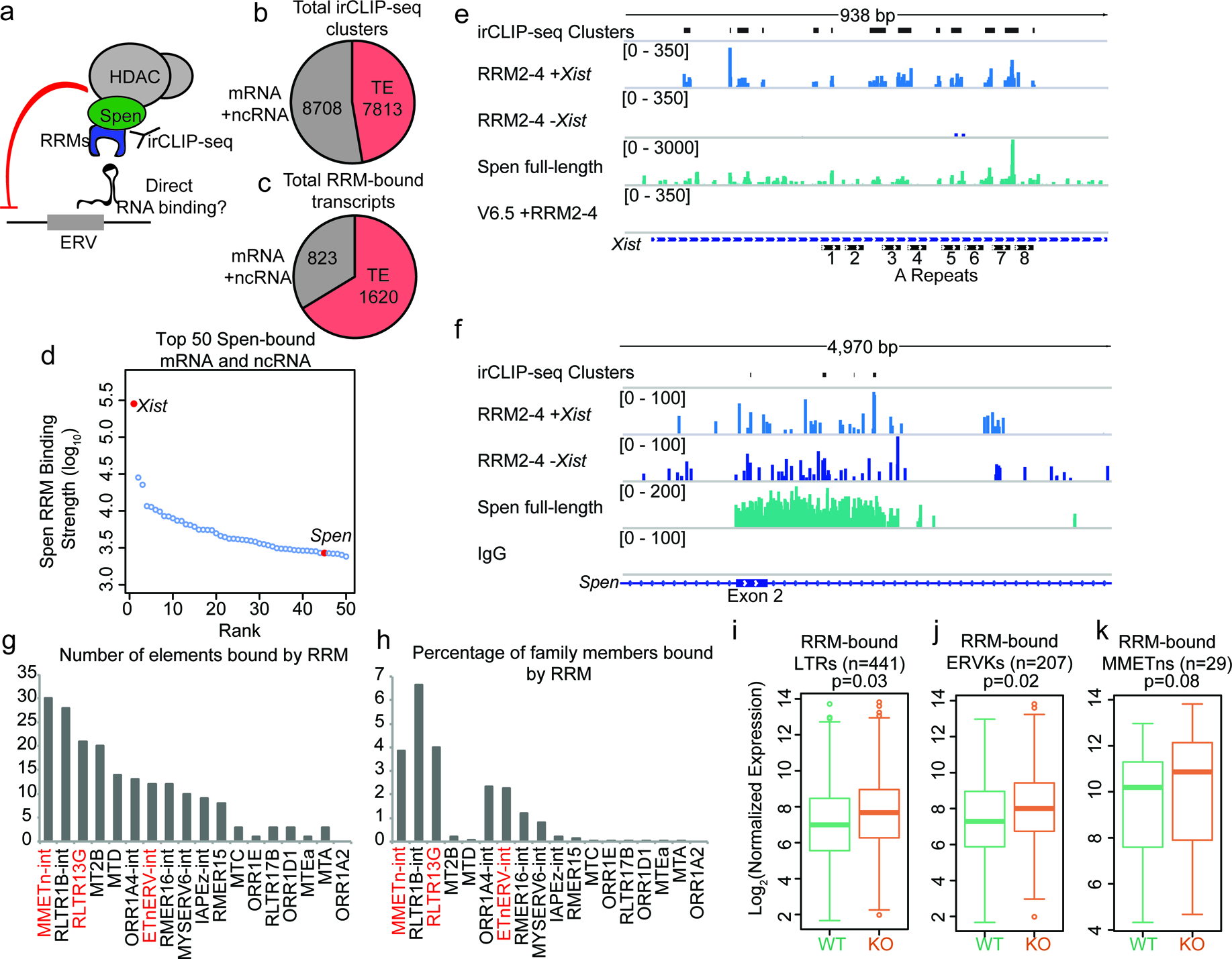
Spen’s RNA binding domains bind specifically to ERVK RNAs *in vivo*. a. Diagram of irCLIP-seq rationale. irCLIP-seq was performed for the Spen RNA binding domains to identify direct RNA binding partners of Spen genome-wide. b. Pie chart showing the number of irCLIP-seq clusteres called on TE RNAs and mRNAs and ncRNAs. c. Pie chart showing the number of RNA transcripts containing at least one irCLIP-seq cluster in TEs and mRNAs and ncRNAs. d. Binding strength of RRM2-4 for the top 50 Spen-bound RNAs. *Xist* (#1) and *Spen* (#40) are highlighted in red. e. Tracks showing the irCLIP-seq signal across the A-repeats of *Xist*. Tracks are shown for Spen KO +RRM2-4 (+/-*Xist*), Spen full-length, and V6.5 male mESCs. irCLIP-seq clusters are shown in black at the top. f. Tracks showing the irCLIP-seq signal across exon 2 of *Spen*. Tracks are shown for Spen KO +RRM2-4 (+/-*Xist*), Spen full-length, and an IgG control. irCLIP-seq clusters are shown in black at the top. g. Graph showing the number of bound TE loci for each TE subfamily with significant Spen RRM2-4 binding. h. Graph showing the percentage of TE loci bound for each TE subfamily with significant Spen RRM2-4 binding. i. Expression of RRM-bound LTRs in WT mESCs (green) and *Spen* KO mESCs (orange). Shown is the log_2_ transformed normalized RNA-seq read counts. For all boxplots, the thick line represents the median, while the box gives the IQR. *P* values are calculated using a two-tailed T test. j. Same as in i for RRM-bound ERVKs. k. Same as in I for RRM-bound MMETns.

Because the balance in the pluripotency transcriptional network is disrupted in *Spen* KO mESCs, we hypothesized that differentiation may be blocked. To test whether *Spen* KO mESCs can spontaneously differentiate when deprived of self-renewal signals, we removed leukemia inhibitory factor (LIF) from the media and allowed cells to grow for 6 days. After 6 days, WT mESCs downregulated pluripotency factors such as Oct4 and expressed early differentiation markers such as Nestin. In contrast, *Spen* KO mESCs maintained Oct4 expression and mESC morphology despite LIF removal (**Extended Data Figure 3b,c**). We then tested whether *Spen* KO mESCs can be directed to differentiate toward the neurectoderm fate, by driving differentiation toward neural progenitor cells. After 14 days of differentiation, WT cells express Nestin and have NPC morphology while *Spen* KO mESCs do not (**Extended Data Figure 3d**). Furthermore, we observed massive cell death during *Spen* KO mESC differentiation, finding that by day 10 of directed differentiation, 100% of cells were no longer viable (**Extended Data Figure 3e**). These results support the results from RNA-seq that suggest that Spen loss leads to a failure to differentiate past the pluripotent state.

We next asked whether Spen represses ERVK loci directly or indirectly. Though we observe changes in chromatin accessibility and covalent histone modifications at these loci when Spen is knocked out, Spen is an RBP that does not bind to chromatin directly. We hypothesized that Spen may recognize ERVK RNAs transcribed at these TE loci and recruit chromatin silencing machinery to them, performing a surveillance role against aberrant transcription of these parasitic elements. Thus if Spen is regulating these ERVK loci directly, we would expect that regulation to be RNA-dependent (**Figure 3a**).

To test this, we performed infrared crosslinking immunoprecipitation followed by sequencing (irCLIP-seq), which allows for identification of direct RBP binding sites on RNA^29^. Because of Spen’s large size (∼400 kDa) and the lack of highly specific antibodies to it, we expressed a FLAG-tagged version of the Spen RNA binding domains (RRM2-4) for irCLIP-seq. We also compared irCLIP-seq data performed using an antibody against the endogenous, full-length Spen protein, despite the higher level of non-specific signal in those experiments^30^. RRMs 2-4 were shown to be *bona fide* RNA binding domains, with RRM3 specifically required for binding to known Spen binding partner SRA RNA *in vitro*^31^. We expressed tagged Spen RRM2-4 in both *Spen* KO mESCs and WT male V6.5 mESCs, which do not express *Xist*, but do express ERVs. Following UV crosslinking of RNA and protein complexes *in vivo*, we isolated RRM2-4-bound RNAs using an anti-FLAG antibody and prepared libraries from the isolated RNA.

On mRNAs and lncRNAs, we detected Spen RRM2-4 binding at 8,708 sites (irCLIP-seq clusters), on a total of 823 transcripts (**Figure 3b,c**). Compared to other RBPs, Spen binds relatively very few RNAs in irCLIP-seq (**Figure 3d**). *Xist* was the top bound gene, with much higher binding strength than other RNAs, consistent with its known interaction with the A-repeat region of *Xist* and role in XCI^3-7^ (**Figure 3d**). Spen RRM2-4 binds specifically to the A-repeat monomers as reported for full-length and truncated Spen (**Figure 3e**; **Extended Data Figure 4**)^18,30,31^. This confirmed that our FLAG-tagged RRMs bind Spen’s RNA targets *in vivo*. In addition to *Xist*, Spen binds to few mRNAs, one of which is the *Spen* mRNA itself (**Figure 3d,f**). This is consistent with our observation that loss of Spen protein leads to upregulation of *Spen* mRNA expression, suggesting that the Spen protein represses its own RNA output in cis.

**Figure 4:**
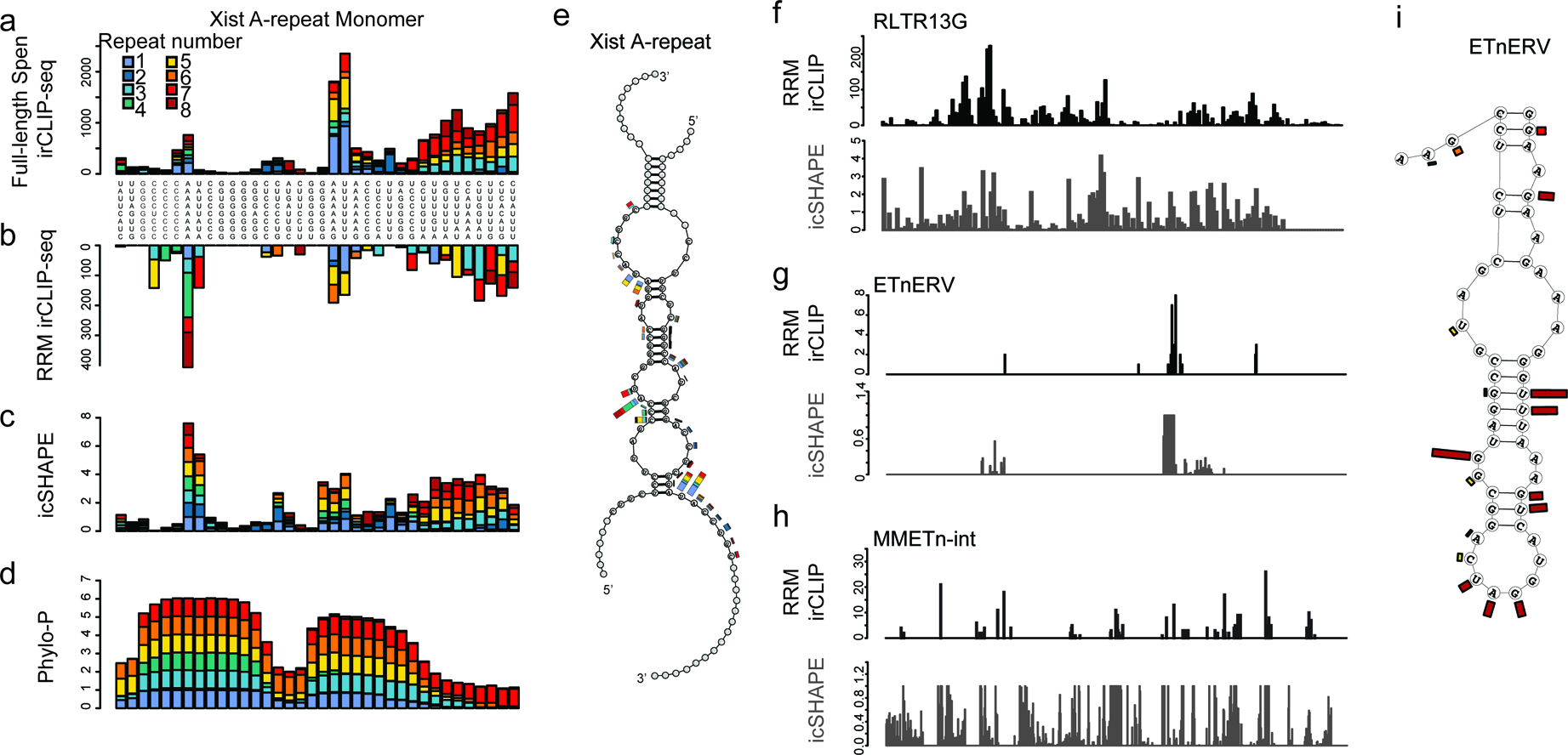
Spen RRMs recognize a common structural feature in A-Repeat and ERVK RNA. a. irCLIP-seq signal from full-length Spen^30^ across the 8 *Xist* A-repeats. Signal for each repeat is stacked across the conserved repeat region. b. Same as in a for RRM2-4 irCLIP-seq. c. icSHAPE reactivity scores for each of the *Xist* A-repeats performed in *vitro*. Signal for each repeat is stacked across the conserved repeat region. d. Phylo-P scores across the A-repeats, showing conservation of repeat sequence at the RRM2-4 binding sites. e. Structural model for Xist A-repeats 4 and 5 which form a hairpin structure. Structural prediction is supported by icSHAPE data. At each nucleotide, the RRM2-4 irCLIP-seq signal strength is plotted as a bar for each repeat (in color). f. irCLIP-seq signal (top, black) and icSHAPE reactivity scores (bottom, gray) for the consensus sequence of RLTR13G elements. Included are elements for which we have both confident RRM2-4 binding and icSHAPE data. g. Same as in f for ETnERV. h. Same as in f for MMETn-int. i. Structural model for ETnERV. Structural prediction is supported by icSHAPE data. At each nucleotide, the RRM2-4 irCLIP-seq signal strength is plotted as a bar.

To assess whether Spen RRM2-4 binds directly to ERVK-derived RNA, we mapped irCLIP-seq data to TEs in the genome. Because irCLIP-seq reads are short and single ended and thus less likely to map uniquely to these repetitive elements, we mapped reads only to the elements that are expressed in our RNA-seq data. We detected a total of 7813 irCLIP-seq clusters on TE RNAs (20% of total clusters detected), covering 1620 individual TEs (**Figure 3b,c**). Of these, we detected RRM2-4 binding at 442 LTRs and 208 ERVKs. The TE subfamily with the greatest number of bound elements is the MMETn-int family, a member of the ETn class (**Figure 3g,h**). In addition, RLTR13G and ETnERV-int showed a large number of elements bound as well as a high percentage of total genomic elements bound (**Figure 3g,h**). RRM-bound elements within these families show modestly increased RNA expression in Spen KO mESCs, consistent with Spen binding contributing to repression of these sites (**Figure 3i-k**). These ERV RNAs are bound by RRM2-4 as well as full-length Spen (**Extended Data Figure 5**).

**Figure 5:**
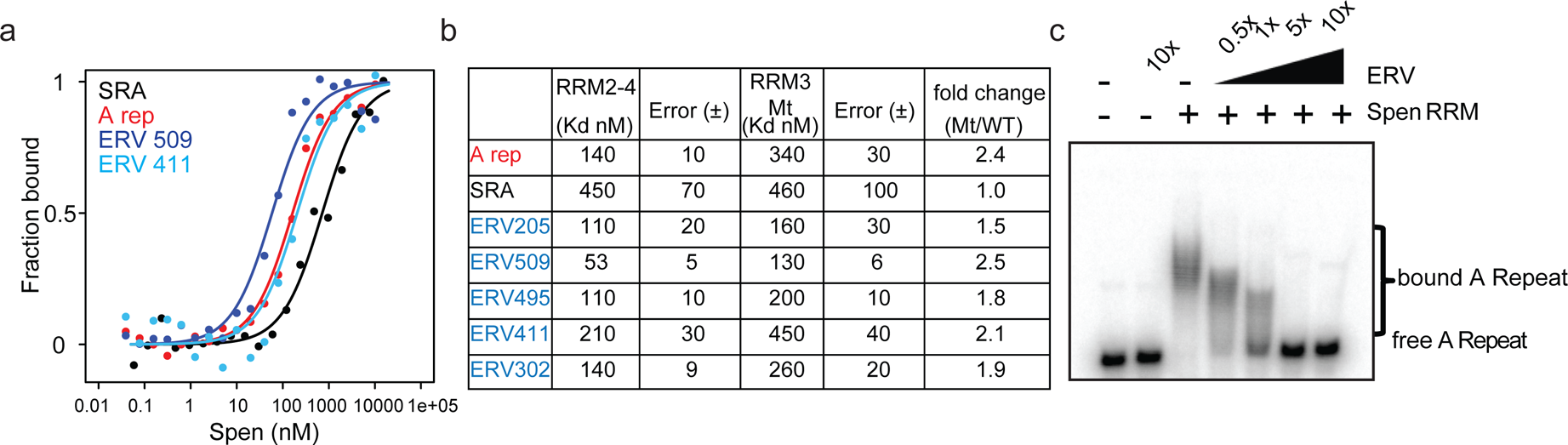
ERVKs bind Spen RRMs in vitro and compete with the A-Repeat. a. Representative binding curves for Spen RRM2-4 from fluorescence anisotropy experiments. Fluoresence anisotropy values are normalized with saturation and offset parameters from the fit to yield fraction bound. Shown are binding curves for *SRA*, the A-Repeat region of *Xist* (A rep), and two *ERV* RNAs bound in irCLIP-seq data (ERV509, ERV411). b. Direct binding constants (K_D_) between RNAs of interest and the Spen RRM2-4 and RRM3 Mt proteins from fluorescence anisotropy experiments. Error is the standard error. The fold change in binding between RRM2-4 and RRM3 Mt for each RNA is shown at the right. c. Acrylamide gel image showing competition of ETn509 RNA with ^32^P-labeled A-Repeat for binding to Spen RRM2-4. Increasing amounts of unlabeled ETn509 competitor RNA (0.5x, 1x, 5x, and 10x the amount of labeled A-Repeat) was added while the protein and A-Repeat concentrations were kept constant. The higher band depicts Spen-bound A-Repeat, while the lower band shows the unbound A-repeat RNA.

The specificity of RBP binding to its RNA targets may have a structural and/or sequence basis. In order to understand whether Spen binds to the A Repeat and ERVK RNAs in the same manner, we first investigated the sequence features in Spen-bound mRNAs, lncRNAs, and ERVKs. We found that RRM2-4 binding sites on mRNA and ncRNAs, as well as ERVKs, are enriched for AU content, but did not find any longer, more complex motifs shared amongst the classes. Furthermore, we did not find sequences that showed strong homology to the highly conserved A Repeat regions (**Extended Data Figure 6**). This was perhaps not surprising, as Arieti *et al.*^31^ previously showed that Spen’s recognition of the *SRA* lncRNA is largely dictated by RNA secondary structure, not sequence, with Spen binding to a flexible, single stranded region adjacent to a double stranded region.

**Figure 6:**
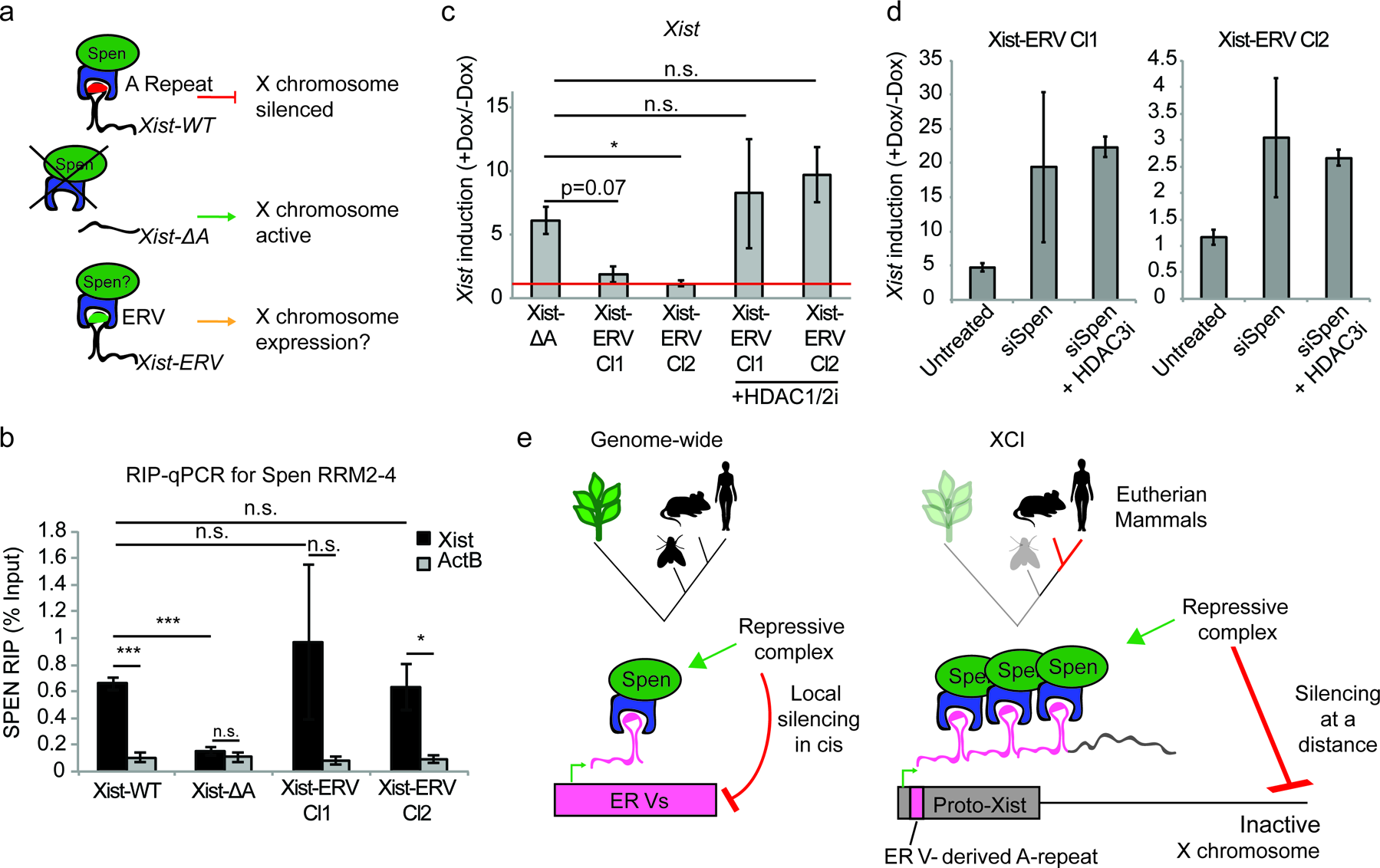
Insertion of an ERV element in place of A-repeat restores Spen binding and local Xist silencing. a. Diagram showing the experimental rationale for replacing the A-Repeat with a 9x tandem insert of an ERV-derived Spen binding site. b. Xist-ERV restores Spen recruitment. RIP-qPCR for *Xist* and *ActB* in Xist-WT, Xist-ΔA, and Xist-ERV clones 1 and 2 mESCs. qRT-PCR values are normalized to input samples. Error bars represent standard error for 5 independent biological replicates. p values are derived from two-tailed *t*-test. c. Xist-ERV causes epigenetic silencing of the *Xist* locus. *Xist* expression level in +dox compared to –dox conditions for Xist-ΔA mESCs and Xist-ERV mESCs with and without HDAC1/2 inhibitor treatment. *Xist* levels are first normalized to *ActB* and then to –dox levels. The red line is drawn at y=0, representing no increase in expression of *Xist* with the addition of doxycycline. For each condition, 4 independent experiments were conducted. p values are derived from two-tailed T test. d. Xist-ERV silencing requires Spen and HDAC3. *Xist* expression level in +dox compared to –dox conditions for Xist-ΔA mESCs and Xist-ERV mESCs with and without siRNA-mediated knockdown of Spen or siRNA-mediated knockdown of Spen + HDAC3 inhibition. For each condition, 4 independent experiments were conducted. e. Model for how Spen evolved to bind *Xist* through its ERV binding mechanism in eutherian mammals.

We asked whether Spen RRMs bind to a similar structural feature in the A Repeat and in ERVK RNAs. Spen has previously been reported to bind to the junction between a single stranded loop and a duplex formed by multiple A-Repeat monomers interacting in 3 dimensions^18^, a structure that is similar to Spen’s binding site within SRA^31^. Upon closer examination of icSHAPE data, which determines whether RNAs are single (high icSHAPE reactivity score) or double stranded (low icSHAPE reactivity score) *in vivo*^32^, we found that the A Repeat region of *Xist* consists of a hairpin that contains a small bulging single stranded region, flanked by two larger single stranded loops (**Figure 4a-e**). Both the Spen RRMs and the full length Spen bind to the conserved single stranded regions directly adjacent to the double-stranded hairpin and to the small bulge within the hairpin (**Figure 4a-e**).

To test whether Spen RRMs bind a similar structural motif in ERVKs, we analyzed icSHAPE data within repeat regions of the genome^33^. For LTR families for which we had enough loci that are both bound by RRMs and have icSHAPE signal, (RLTR13G, ETnERV, MMETn) we found that Spen binding occurs within short single stranded regions adjacent to double stranded regions (**Figure 4f-h**). This is consistent with a model where Spen RRMs bind to single-stranded bulges within larger duplexes in ERVK RNAs (**Figure 4i**; **Extended Data Figure 7a,b**). Taken together, this demonstrates that Spen RRMs bind to the A-repeat of Xist and to ERVK RNAs in the same manner, recognizing a specific structural feature in the RNA.

Though irCLIP-seq provides a transcriptome-wide map of Spen RRM-RNA binding, we wanted to understand in even more depth the specificity and strength of the interaction between the Spen RRMs and *ERVK* RNAs. We hypothesized that if Spen binds *ERVK*s and the A Repeat with the same structural recognition mechanism, it should bind with similar affinity and in a competitive manner. We used fluorescence anisotropy experiments to measure the binding affinity of Spen RRMs 2-4 to *ERV* RNA elements as well as the *Xist* A-repeat and the *SRA* RNA H12H13 hairpin, two known interactors of Spen^3,4,18,31^. We also measured binding of a version of the Spen RRMs that has 5 point mutations in RRM3 (RRM3 Mt), which was previously shown to diminish binding of Spen to SRA (**Extended Data Figure 8a**)^31^. Using *in vitro* irCLIP RNA isolation, we found that the RRM3 Mt binds ∼50% less RNA than RRM2-4 confirming these residues are essential for Spen RNA binding (**Extended Data Figure 8b-f**).

Using fluorescence anisotropy, we found that the binding affinity of Spen RRM2-4 for our *ERV* transcripts and the *Xist* A-Repeat are similar (dissociation constant *K_D_* = 53nM ∼140 nM), and higher than the affinity of Spen RRM2-4 for the *SRA* hairpin (K_D_ = 450 nM) (**Figure 5a,b**). The binding affinity of the RRM3 Mt is about half that of the RRM2-4 for both *Xist* A-repeat and the *ERV* RNAs, suggesting that the same residues of Spen’s RRM3 domain are binding both *Xist* A-repeat and *ERV* RNAs (**Figure 5b**; **Extended Data Figure 8g**). Finally, we found that an unlabeled *ERV* RNA competed with a labeled *Xist* A-repeat for binding to Spen RRM2-4, further demonstrating that Spen recognizes these two RNAs, in part, through the same binding domain and in a very similar manner (**Figure 5c**). These data, taken with our irCLIP-seq data, show that Spen binds to ERVK-derived RNA in the nucleus and does so via the same binding mechanism and with the same strength as it binds the A-Repeat of *Xist*. This supports a model in which Spen directly represses ERVK loci in the genome by recognizing their RNA transcripts through their secondary structures.

Collectively, these results show that Spen’s RNA binding domains bind TE-derived RNA in mESCs and do so via the same mechanism that they bind *Xist* RNA, with a significant contribution from the RRM3 domain. The observation that the A-repeat may be derived from an ancient ERV insertion^9^, which shows structural similarity to the ERVK elements bound by Spen, suggests that *Xist* evolved its ability to recruit Spen via Spen’s recognition of ERV sequence and subsequent recruitment of repressive complexes. Spen itself has evolved relatively little throughout evolution from fish to humans, and Spen’s RNA binding domains show no evidence of branch-specific evolution in the eutherian clade (**Extended Data Figure 9a-h**). Thus a mechanism in which the *Xist* RNA itself evolved toward Spen’s functionality is fitting.

We reasoned that if the *Xist* A-repeat region evolved via the insertion of a TE that had affinity for Spen into the proto-*Xist* locus, then the insertion of an ERV element into *Xist* may be able to complement the A-repeat deletion of *Xist*. We would expect that this chimeric *Xist-ERV* would be able to 1) recruit and bind to Spen and 2) silence X-linked genes in *cis* (**Figure 6a**). To test this hypothesis, we utilized a male (XY) mESC line that harbors both a doxycycline inducible *Xist* gene on the X as well as a deletion of the A-repeat region (Xist-ΔA). While induction of *Xist* by doxycycline treatment in the WT male mESCs (Xist-WT) leads to silencing of the single X chromosome, *Xist* expression in Xist-ΔA mESCs has no effect on X chromosome gene expression or accessibility^8,10^. We used CRISPR-Cas9 genome editing to insert a 9x array of a 154-bp ERV-derived Spen binding site from chromosome 16 into the X chromosome where the A-repeat has been deleted in Xist-ΔA mESCs (Xist-ERV) (**Extended Data Figure 10a-c**). We chose a 9x array of binding sites because the A-repeat region of *Xist* consists of 8.5 repeats in human and 7.5 repeats in mouse, which fold into a complex secondary structure with multiple Spen binding sites^18,34^. Furthermore, the addition of monomers of the A-repeat up to 9 repeats in an A-repeat knockout background leads to a linear increase in chromosome silencing capacity^8,34^.

We first asked whether *Xist-ERV* is able to bind directly to and recruit Spen. To do this, we expressed the FLAG-tagged Spen RRM2-4 in our cell lines containing Xist-WT, Xist-ΔA, or Xist-ERV transgenes (2 independent clones), and measured *Xist*-RRM binding under optimized expression conditions (more below), by RIP-qPCR and CLIP-qPCR. We found that *Xist-ERV* is bound by Spen RRM2-4, similar to *Xist-WT* and in contrast to *Xist-ΔA* which could not bind Spen (**Figure 6b**; **Extended Data Figure 11a,b**). Next, when we quantified the ability to induce *Xist-ERV* expression with the addition of doxycycline, we were surprised to find that expression of *Xist-ERV* could not be induced, in comparison to *Xist-WT* and *Xist-ΔA* (**Figure 6c**). Genetic analysis confirmed that the Tet operator and CMV promoter were intact in all of our *Xist-ERV* clones (**Extended Data Figure 10d**), suggesting that the lack of gene induction may be due to epigenetic silencing recruited by Spen. To test whether the inability to induce *Xist-ERV* expression was due to silencing in *cis*, we treated cells with an HDAC1/2 inhibitor (HDAC1/2i) or an HDAC3 inhibitor (HDAC3i) for 24 hours before *Xist* induction. We found that both HDAC1/2i and HDAC3i de-repressed the transgene and *Xist-ERV* was expressed (**Figure 6c**; **Extended Data Figure 10e-g**).

To test whether this *Xist-ERV* silencing was Spen-dependent, we used siRNAs to knockdown Spen 24 hours prior to induction of *Xist* with doxycycline. We found that *Xist-ERV* expression was elevated in Xist-ERV clones 1 and 2 after Spen knockdown (**Figure 6d**). The rescue of *Xist-ERV* expression by Spen depletion was quite variable from experiment to experiment; addition of an inhibitor of HDAC3, which is recruited by Spen^2,4^, made the Xist-ERV induction substantially more constitent and up to 20-fold in at least one clone (**Figure 6d**). This suggested that *Xist-ERV* recruits Spen, leading to local silencing of the *Xist* transgene in *cis*.

Therefore, the insertion of an ERV-derived RNA motif into *Xist* is sufficient to partially substitute for the A-repeat, recruit Spen to the chimeric RNA, and mediate local gene silencing *in cis*. This is the first demonstration of how a TE insertion during evolution could confer new functionality on a noncoding RNA, by introducing a new protein recruitment domain^11^.

Finally, we tested whether this Spen recruitment by the *Xist-ERV* RNA is sufficient to lead to chromosome-wide silencing. We conducted these experiments in the presence of HDAC1/2 inhibition, which increases *Xist* induction but was previously shown not to affect silencing during XCI^2^. While expression of X-linked genes *Pgk1*, *Gpc4*, *Mecp2*, and *Rnf12* was reduced after *Xist* induction in Xist-WT cells, expression was unchanged upon expression of *Xist*-*ΔA* or *Xist-ERV* (**Extended Data Figure 11c**). Thus, while the addition of the ETn array to an A-repeat-deficient *Xist* was sufficient to recruit Spen, it was not sufficient to carry out XCI. These experiments were conducted in the presence of HDAC1/2i, necessary to overcome Xist-ERV silencing of its own locus, but may also inhibit the chimeric RNA’s silencing of distal loci. Furthermore, this could be due to the differences in levels of *Xist* that we are able to induce in this system, which are lower in the Xist-ERV cells than the Xist WT or Xist ΔA cells, even in the presence of HDAC1/2i. Nevertheless, our results are consistent with the hypothesis that insertion of an ERV element into the proto-*Xist* locus was an *early* evolutionary step in the functionalization of the A-repeat, but that additional rounds of evolution may have subsequently acted on this region of the genome in order to evolve *Xist*’s full functionality for silencing the entire chromosome over long genomic distance^9,35^.

## DISCUSSION

Here we show that Spen, a highly conserved and pleiotropic RNA-binding protein, binds to and regulates specific classes of ERVK loci in mouse embryonic stem cells. Spen represses these ERVK loci via a direct mechanism where it binds transcripts derived from these parasitic elements, and can then recruit its chromatin silencing protein partners. In the absence of Spen, these elements become derepressed, showing loss of repressive H3K9me3 and gain of active chromatin modifications. Sensing transposon silencing at the RNA level is attractive, as it provides a mechanism for the cell to identify the “leaky” locus and target the silencing machinery to the locus in *cis*.

These observations are striking when taken with the observation that the *Xist* RNA, which is also robustly bound by Spen, is derived from a number of ancient retroviral insertions^9^. We show direct evidence, for the first time, that an *Xist* binding protein can bind to ERV-derived RNA *in vivo*, and thus may have been recruited to *Xist* via the insertion of an ERV element into the proto-*Xist* locus (**Figure 6e**). This evidence supports the hypothesis that lncRNAs, which are evolutionarily young, may exapt TE-protein interactions in order to form functional protein binding domains^11,12,36,37^. In effect, *Xist* lncRNA “tricks” the female cell to perceive the inactive X chromosome as being coated by dangerous transposon RNAs. The cell can then deploy Spen, a transposon-silencing protein, to carry out X chromosome dosage compensation.

One of the major mysteries of X inactivation is the *cis*-restriction of *Xist*, which only silences the chromosome from which it is expressed, a property critical for dosage compensation of two X chromosomes in females to one X in males. Our discovery of the link between Spen and transposon silencing suggests an ancient evolutionary origin for *cis* silencing. Spen sensing of *ETn* RNA and Spen-mediated silencing are *cis* restricted, and indeed grafting ETn into the *Xist* locus caused *cis* silencing of *Xist* RNA expression itself. Hence, these data suggest that the invasion of an ERV-like sequence into proto-*Xist* led first to RNA-based silencing in *cis*. Subsequent mutations in this proto-*Xist* may then have allowed this RNA to localize to and silence distal sites on the X chromosome. The transition from short to long range silencing in *cis* may be a fruitful topic in future studies.

## Contributions

A.C.C. and H.Y.C. devised the project, designed figures, and wrote the manuscript. A.C.C. performed all experiments except for *in vitro* biochemistry. J.X. and A.C.C. performed sequencing data analysis. M.Y.N., R.T.B., and D.S.W. designed and performed *in vitro* biochemistry experiments. Y.W. performed irCLIP-seq analyses. Q.S. helped with cloning of constructs. R.L. helped with RNA preparations. K.E.Y. and D.R.D. helped with cell line maintenance. J.P.B. helped with design of irCLIP-seq experiments. R.C.R., A.S., and S.N. FACS sorted cells. S.N. and S.W.C. helped with plasmid design and cloning strategies. A.M. provided recombinant proteins for use in irCLIP-seq experiments. All authors contributed to the writing of the manuscript.

## Acknowledgments

We thank E. Heard and A. Wutz for reagents and for helpful discussions. We thank F. Fazal, M.R. Corces and members of the Chang lab for assistance and feedback. Supported by NIH RM1-HG007735, R01-HG004361, and Scleroderma Research Foundation (to H.Y.C.), R01-GM120347 (to D.S.W., R.T.B.), and the Hagey Laboratory for Pediatric Regenerative Medicine and R01DEO26730 (to M.T.L). H.Y.C. is an Investigator of the Howard Hughes Medical Institute.

## Data Availability Statement

All raw and processed sequencing data can be found in GEO under accession number GSE131413.

## Code Availability Statement

Analysis methods used for next generation sequencing data in this paper are described in the appropriate sections of the methods. Any code used in this study will be made available by the authors upon request.

---

**Extended Data Figure 1:**
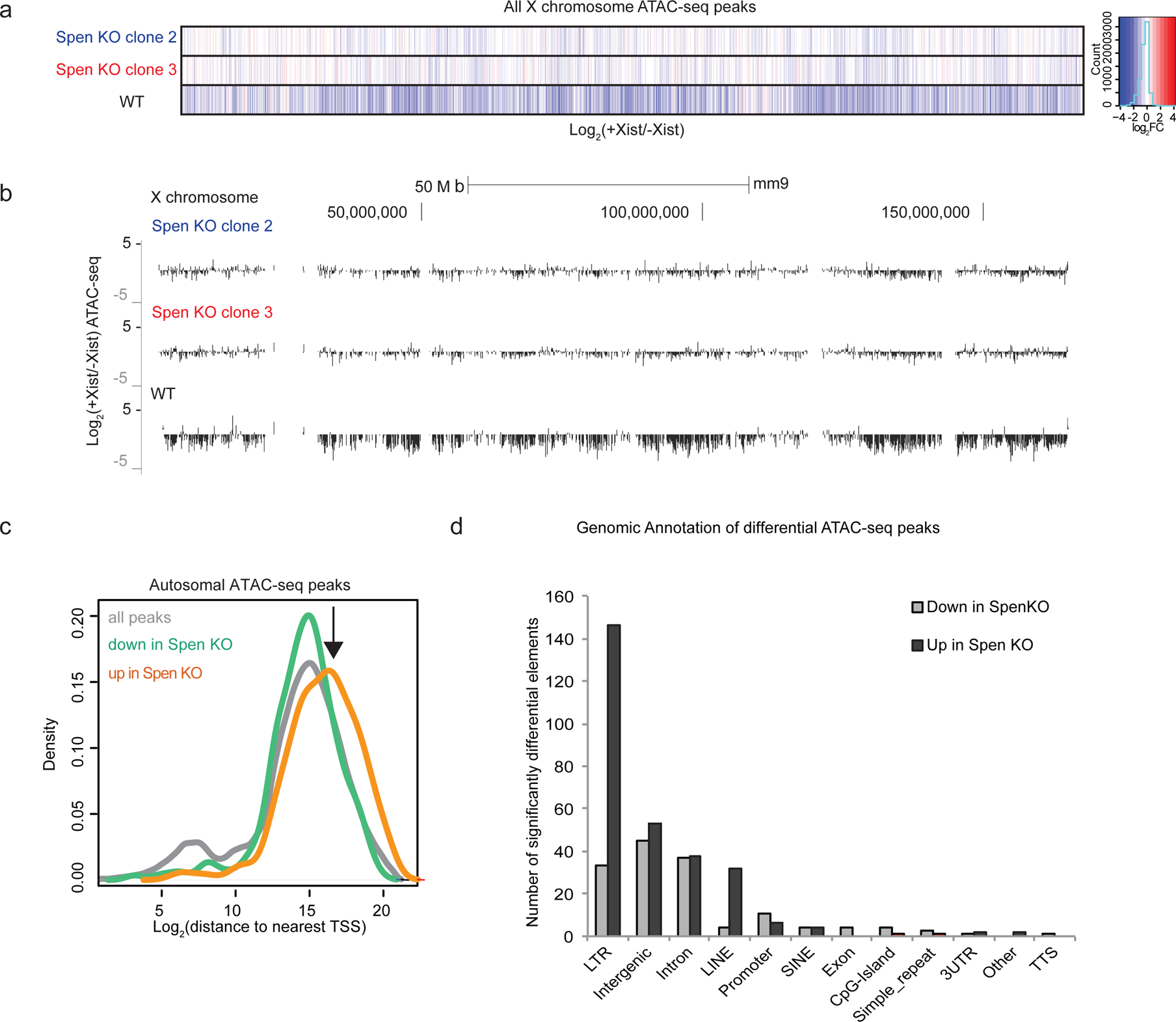
Spen is required for XCI chromosome-wide. a. Heatmap showing the log_2_(fold change) in +Dox/-Dox conditions in WT and *Spen* KO clones 2 and 3 for all ATAC-seq peaks on the X chromosome. b. Tracks showing the Log_2_ ratio of ATAC-seq signal in +Dox and –Dox samples for 2 *Spen* KO clones and one WT clone after 48 hours *Xist* induction. The entire X chromosome is shown to scale. c. The Log_2_ distance to the nearest coding gene transcription start site (TSS) for all ATAC-seq peaks on the autosomes. All peaks are shown in gray and show a small peak at TSSs and a larger peak at distal sites. Peaks that lose accessibility in *Spen* KO (green) show a similar pattern. Peaks that gain accessibility (orange) in *Spen* KO are distal to TSSs, as indicated by arrow (p-value =5.97×10^-13^ by KS test) and do not overlap promoters. d. Plot showing the number of ATAC-seq peaks covering each genomic annotation from HOMER. Shown are the significantly differential peaks between WT and *Spen* KO mESCs that gain (black) or lose (gray) accessibility.

**Extended Data Figure 2:**
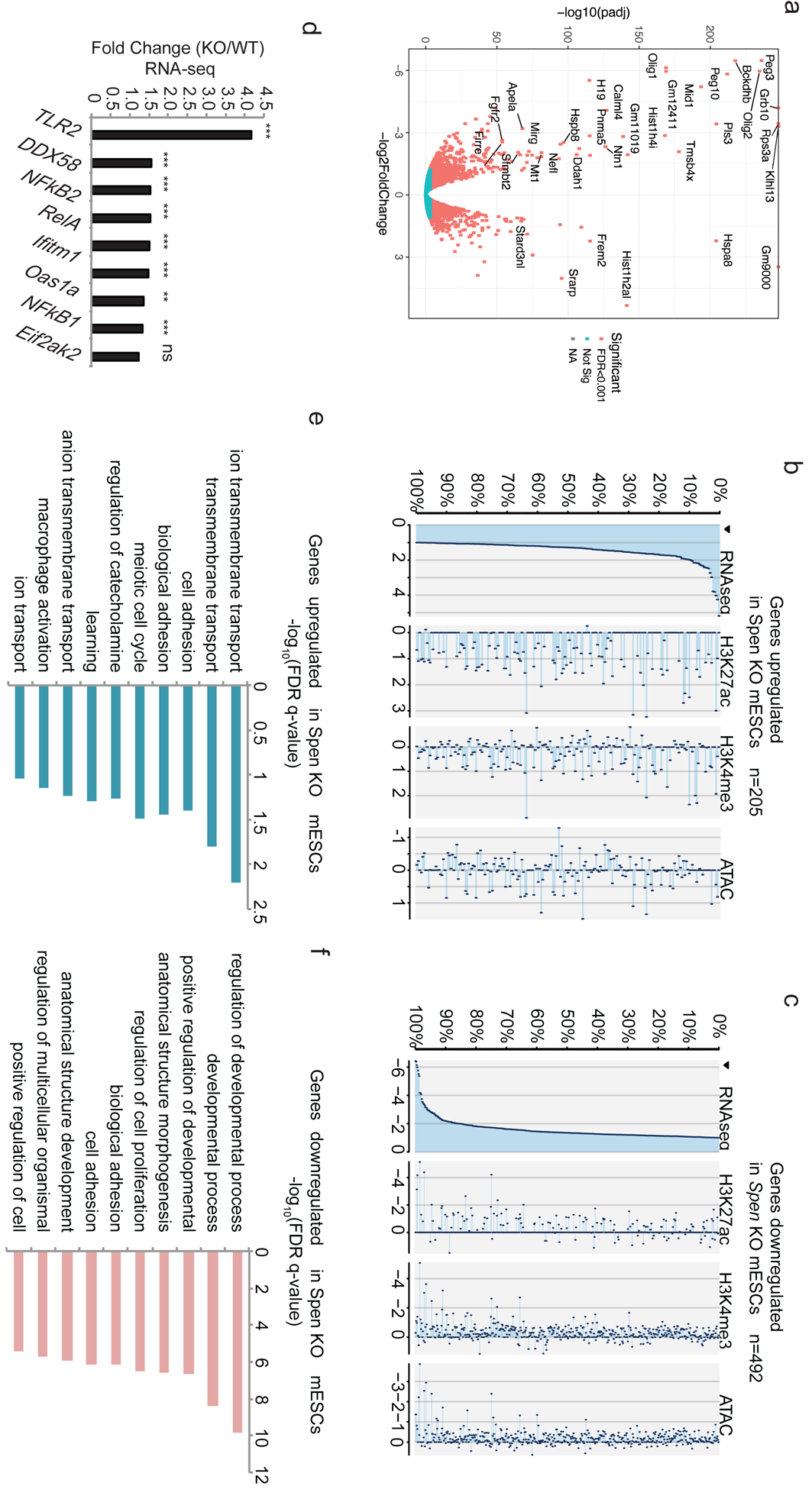
RNA-seq reveals downregulation of developmental genes in *Spen* KO mESCs. a. Volcano plot showing the log_2_ fold change for gene expression (RNA-seq) in WT vs. *Spen* KO mESCs. Genes with negative values are upregulated in *Spen* KO mESCs compared to WT controls. The y-axis shows the –log_10_ (p-adjusted) calculated from DESeq2. b. Histone modifications and chromatin accessibility changes at the promoters of genes upregulated in *Spen* KO mESCs (*n*=205 genes). Plotted is the log_2_ fold change for each mark at the promoter of genes upregulated in *Spen* KO mESCs. c. Same as in b for genes downregulated in *Spen* KO mESCs (*n*=492 genes). d. Fold change (*Spen* KO/WT) in expression (from RNA-seq) for genes involved in the innate cellular immunity pathway. e. GO terms for cellular processes associated with the genes upregulated in *Spen* KO mESCs. GO terms are derived from GORilla and the FDR q-value is calculated from a Benjamini-Hochberg correction of the p-value for enrichment using a ranked list of the genes changing in *Spen* KO. f. GO terms for cellular processes associated with the genes downregulated in *Spen* KO mESCs. GO terms are derived from GORilla and the FDR q-value is calculated from a Benjamini-Hochberg correction of the p-value for enrichment using a ranked list of the genes changing in *Spen* KO.

**Extended Data Figure 3:**
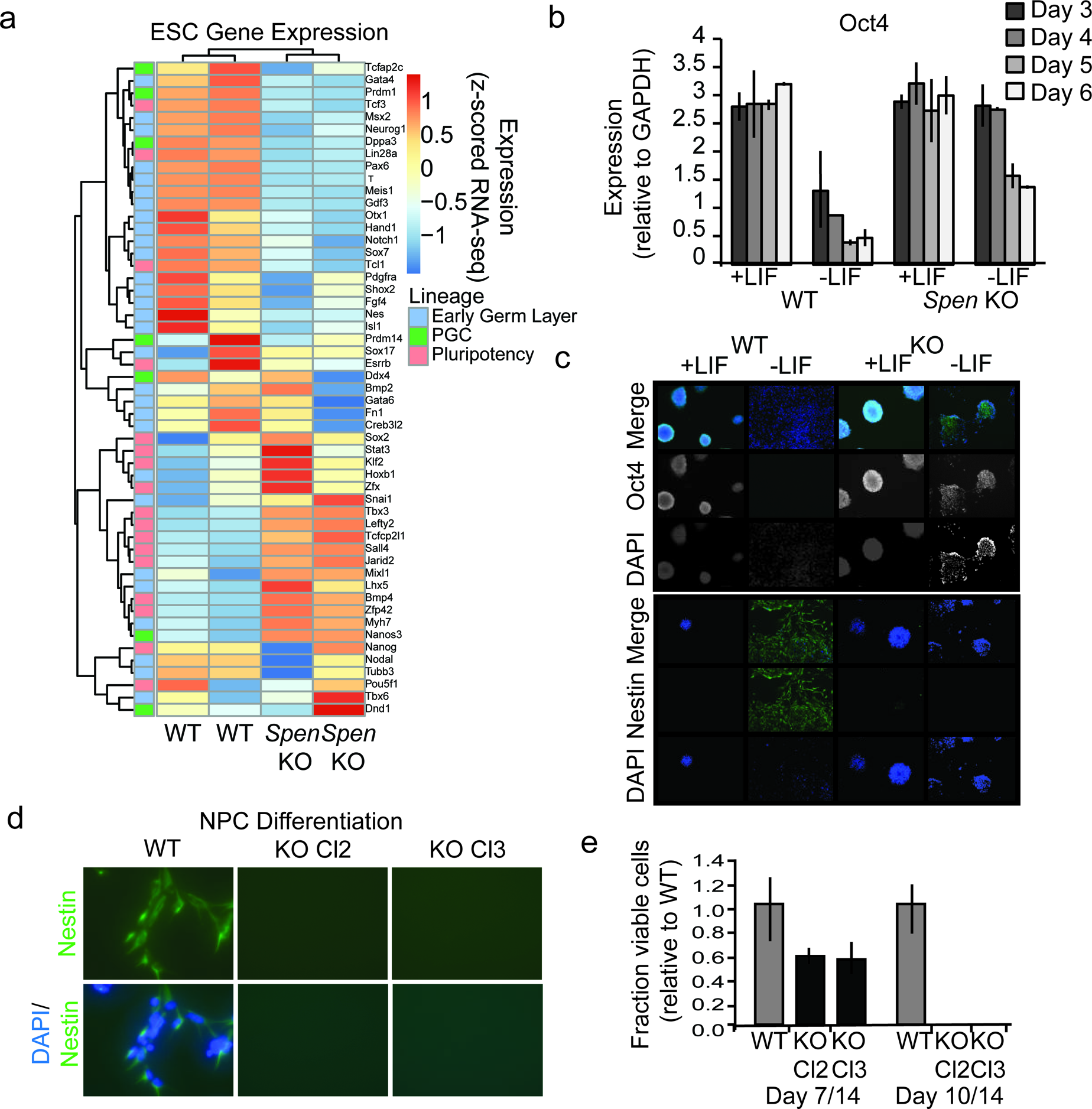
*Spen* KO mESCs fail to differentiate from the pluripotent state. a. Expression of canonical mESC genes in *Spen* KO and WT mESCs. Color on the left indicates whether the genes specify early germ layers (blue), primordial germ cells (green), or self-renewal (pluripotency; pink). Color in the heatmap represents the z-score for normalized expression by RNA-seq in WT and *Spen* KO mESCs. b. *Oct4* expression, measured by qRT-PCR in *Spen* KO and WT cells with or without LIF for 3,4,5, and 6 days. Expression relative to GAPDH is plotted. Error bars represent the standard deviation across 3 independent replicates. c. Representative immunofluorescence images of WT and *Spen* KO mESCs in +LIF and – LIF conditions stained with antibodies against Oct4, and Nestin. DAPI staining shows nuclei. LIF was removed for 4 days, and cells were allowed to differentiate spontaneously *in vitro* before fixing and staining. d. Representative immunofluorescence images of WT and *Spen* KO mESCs stained with anti-Nestin antibody and DAPI after 14 days of *in vitro* differentiation toward neural progenitor cells. Images are acquired at 40x magnification. e. Quantification of the fraction of viable cells (relative to WT) at day 7 and day 14 of NPC differentiation.

**Extended Data Figure 4:**
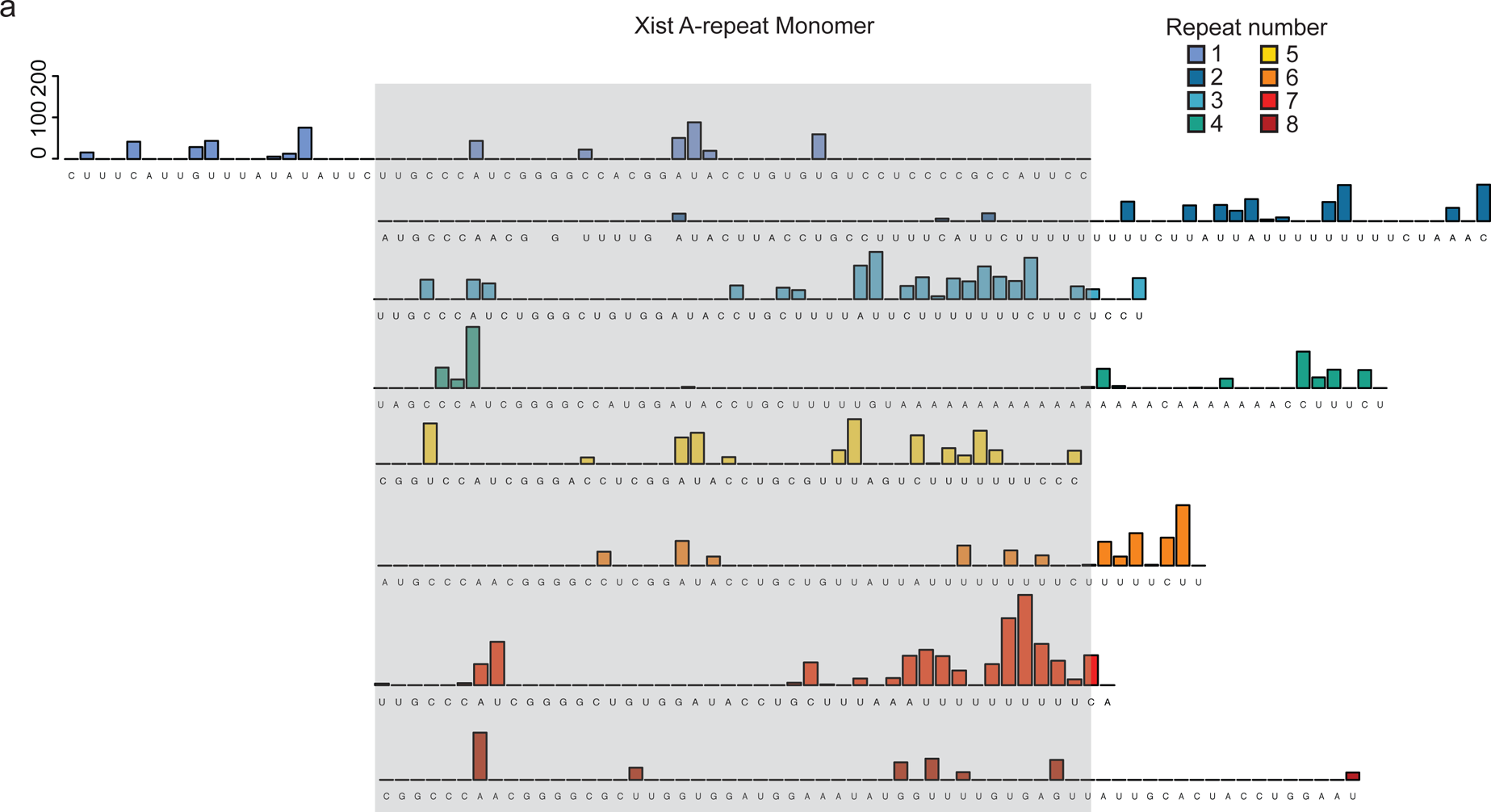
irCLIP-seq of RRM2-4 at the *Xist* A-repeats. a. Related to Figure 4b. irCLIP-seq signal from RRM2-4 shown for each of the A-repeats in mouse. The conserved portion of the repeat is shown stacked together (in gray box) with the linker regions outside.

**Extended Data Figure 5:**
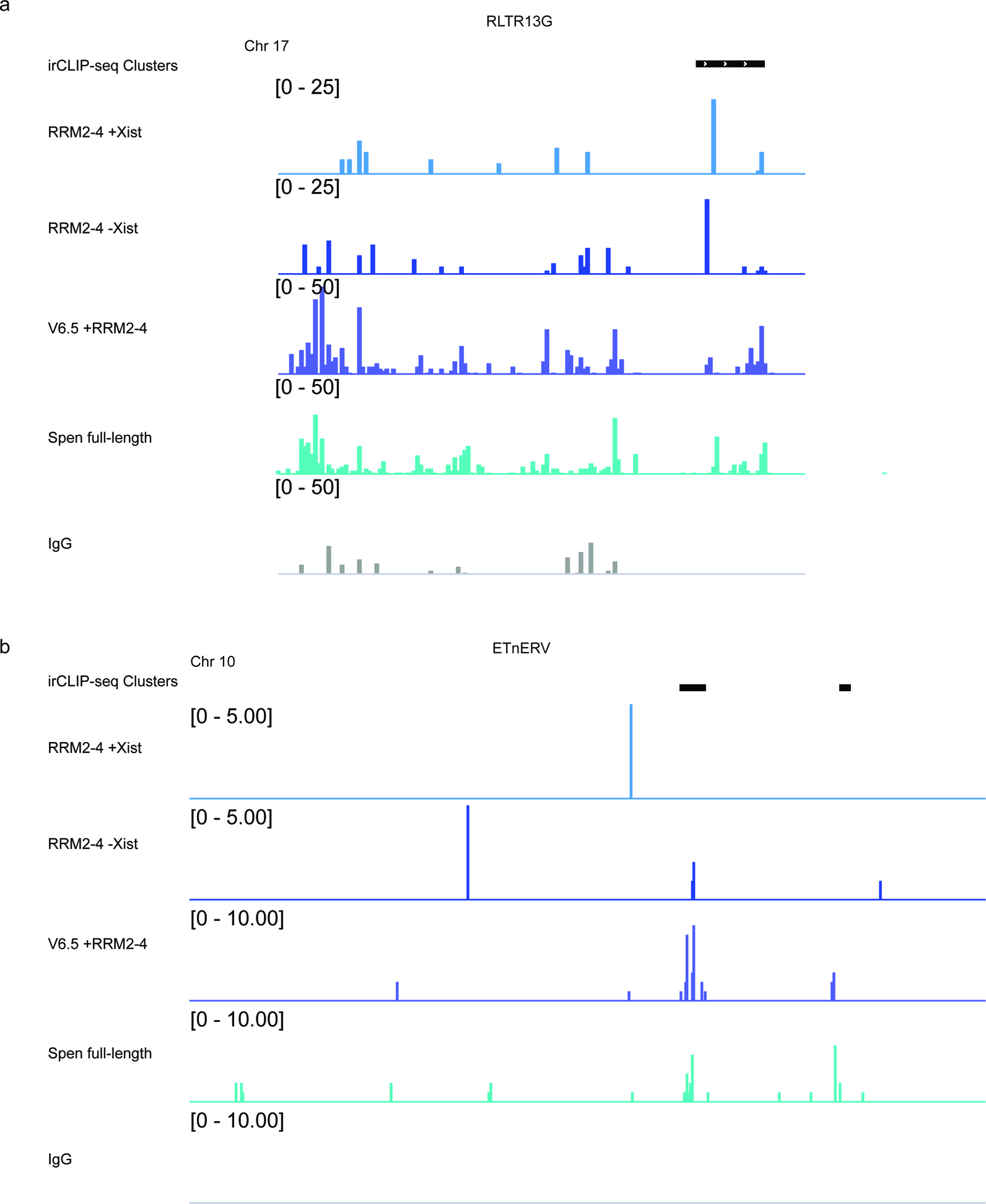
Comparison of RRM2-4 and Spen full length binding to ERVs. a. Tracks showing irCLIP-seq signal for RRM2-4 and Spen full-length at an RLTR13G element on chromosome 17. irCLIP-seq clusters are indicated in black at the top. b. Same as in a for an ETnERV element on chromosome 10.

**Extended Data Figure 6:**
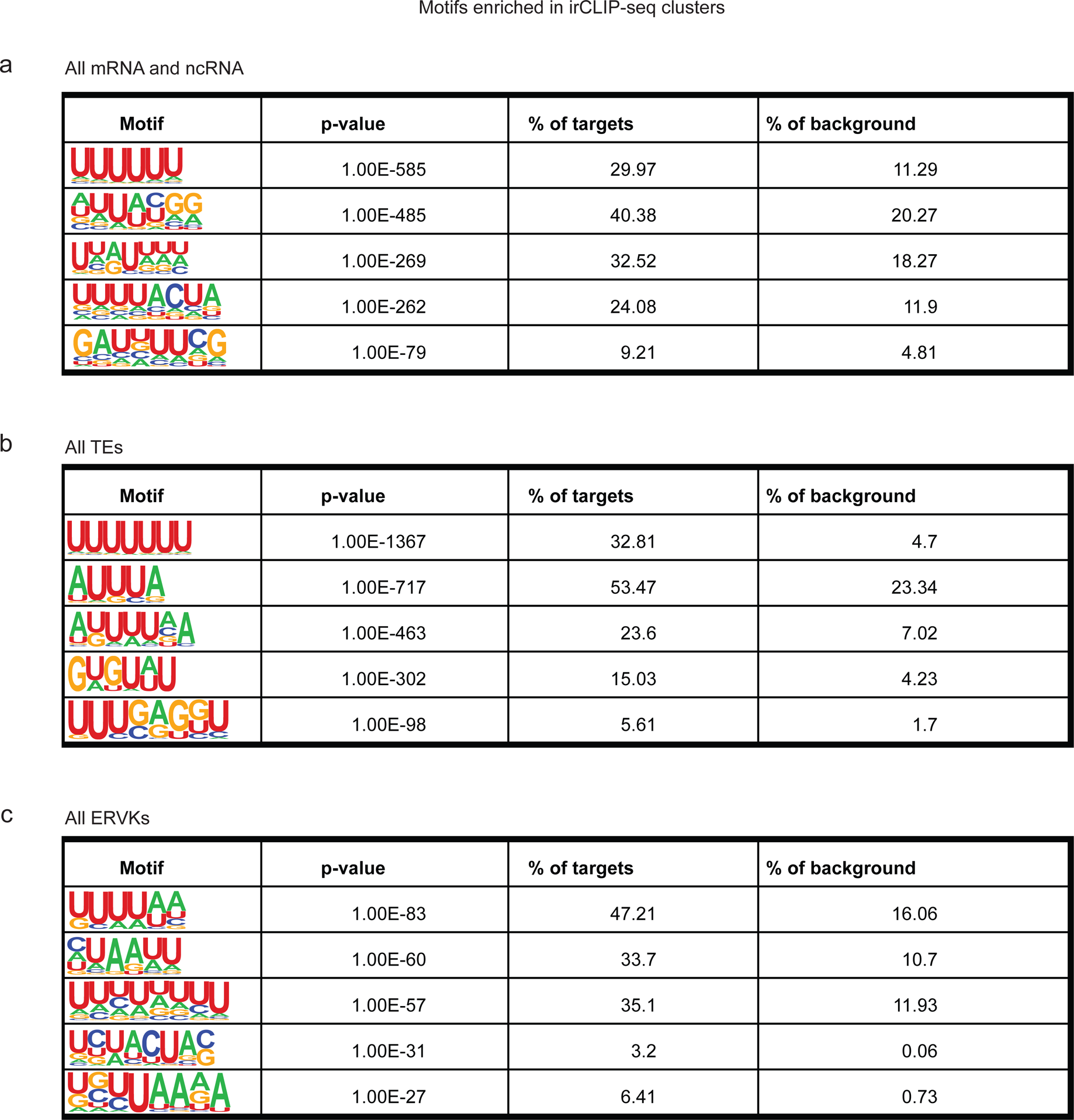
Motif analysis for RRM2-4 RNA binding sites. a. Motifs identified in irCLIP-seq clusters (extened to 50 bp total length) by HOMER for mRNAs and ncRNAs. Presented is the motif, the p-value, the percentage of peaks containing the motif, and the percentage of background sites containing the motif. b. Same as in a for irCLIP-seq clusters in TEs. c. Same as in a for irCLIP-seq clusters in ERVKs.

**Extended Data Figure 7:**
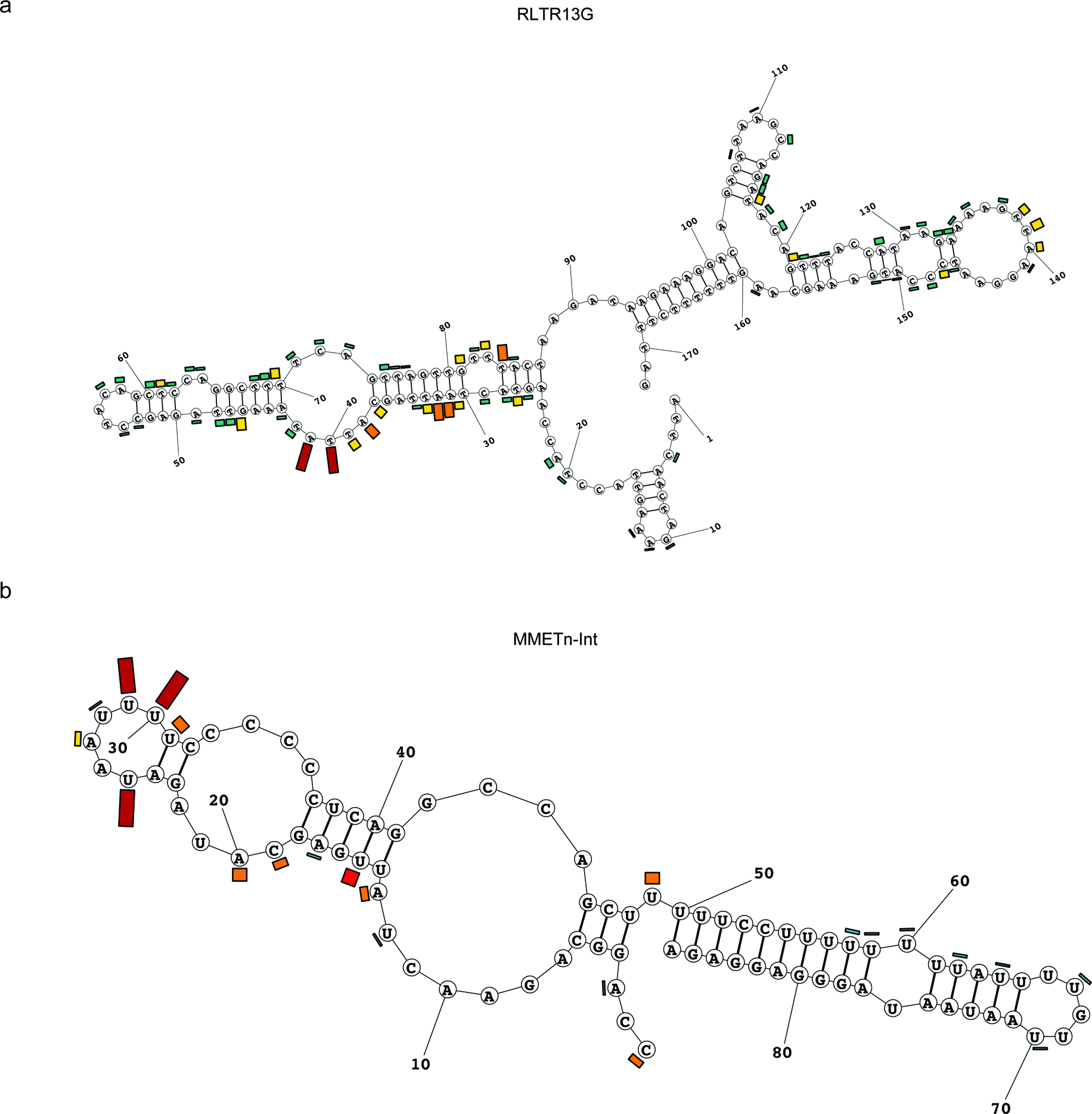
Structural models for Spen-bound ERV RNAs. a. Same as in Figure 4i for an RLTR13G element. b. Same as in Figure 4i for an MMETn element.

**Extended Data Figure 8:**
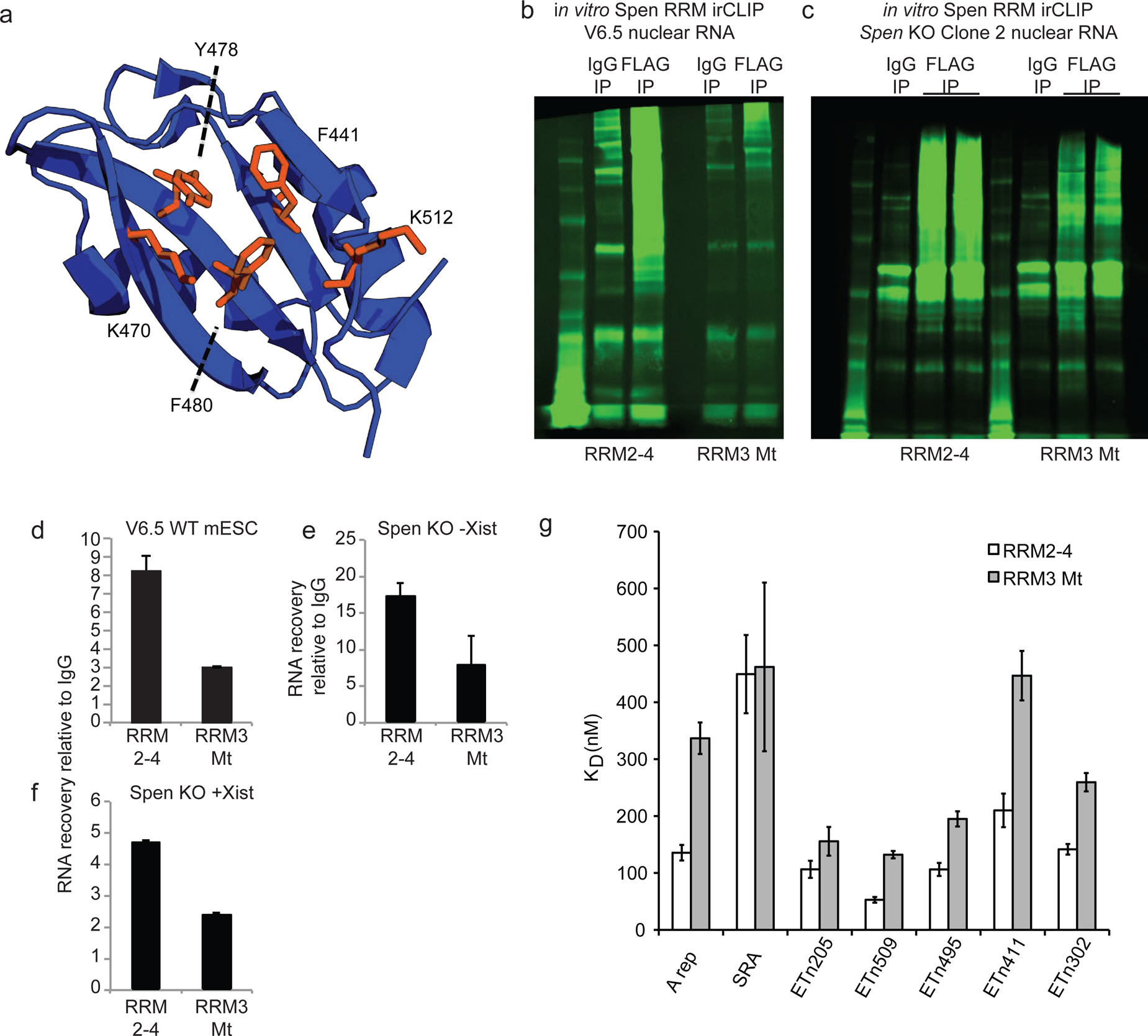
Characterization of RRM3 Mt protein. a. PyMol structure of the RRM3 domain of Spen with the 5 mutated residues indicated. These 5 residues were identified by Arieti et al. as residues important for RNA binding^31^. Figure adapted from PDB: 4P6Q. b. Representative infrared gel image showing ir-labeled RNA after pulldown of RNA-protein crosslinked protein with anti-IgG or anti-FLAG antibodies from V6.5 WT mESCs. FLAG antibody recognizes the Spen RRM2-4-FLAG or Spen RRM3Mt-FLAG protein. The ladder (lefthand lane for each sample) ranges from 12-225 kDa. c. Same as in b for RNA from *Spen* KO cell lines. d. Quantification of the gel in b and an additional replicate. Plotted is the signal intensity in the FLAG lane compared to the IgG lane. Error bars represent the standard deviation from 2 independent biological experiments. e. Same as in d for the gel in c (left). f. Same as in d for the gel in c (right). g. K_D_ values from fluorescence anisotropy experiments for RRM2-4 and RRM3 Mt binding to *SRA* RNA, *Xist* A-Repeat RNA, and *ETn* RNAs (ETn 205, 509, 495, 411, 302). Error bars represent standard error.

**Extended Data Figure 9:**
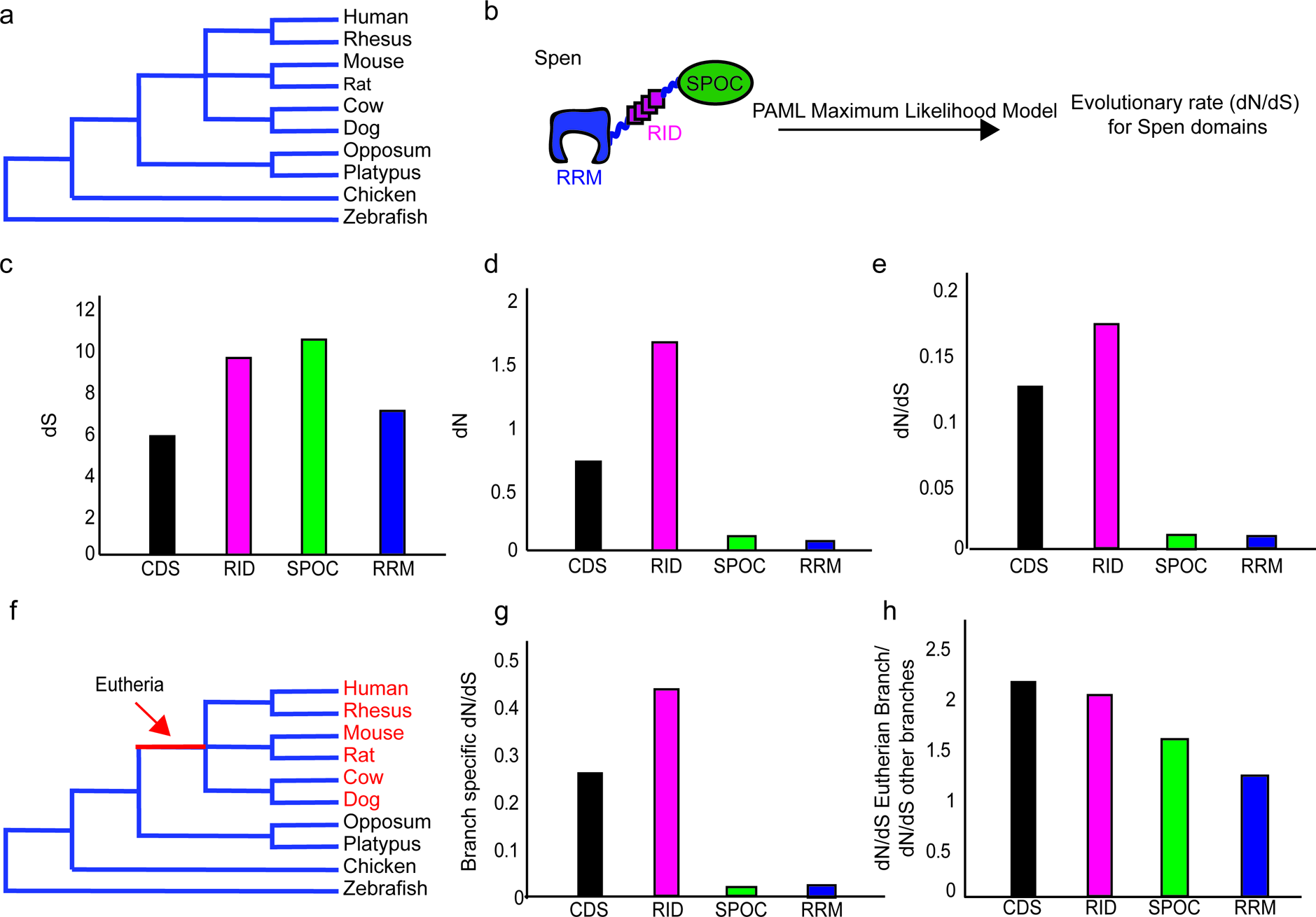
Spen’s RRM domains have not evolved specifically in the Eutherian lineage. a. Diagram of the phylogenetic relationship (not to scale) of eutherian mammals, non-eutherian mammals, chicken, and zebrafish. b. Diagram showing how the evolutionary rate (dN/dS), which shows the ratio of non-synonymous to synonymous changes in the protein, is calculated using a PAML maximum likelihood model for each of the Spen protein domains including the RNA binding domains (RRM), Receptor interaction domain (RID), and protein-interaction domain (SPOC). For each set, the entire coding sequence (CDS) is also included. c. dS for the four illustrated Spen domains during evolution from zebrafish to human. d. dN for the four illustrated Spen domains during evolution from zebrafish to human. e. dN/dS for the four illustrated Spen domains during evolution from zebrafish to human. f. Diagram showing the eutherian branch which is used to calculate whether there is branch-specific evolution in the Spen protein in eutherian mammals. g. Branch specific dN/dS for the eutherian branch for the four illustrated domains of Spen. h. Ratio of the evolutionary rate in the eutherian branch to the evolutionary rate of the other branches. The RRM does not evolve more quickly in this line.

**Extended Data Figure 10:**
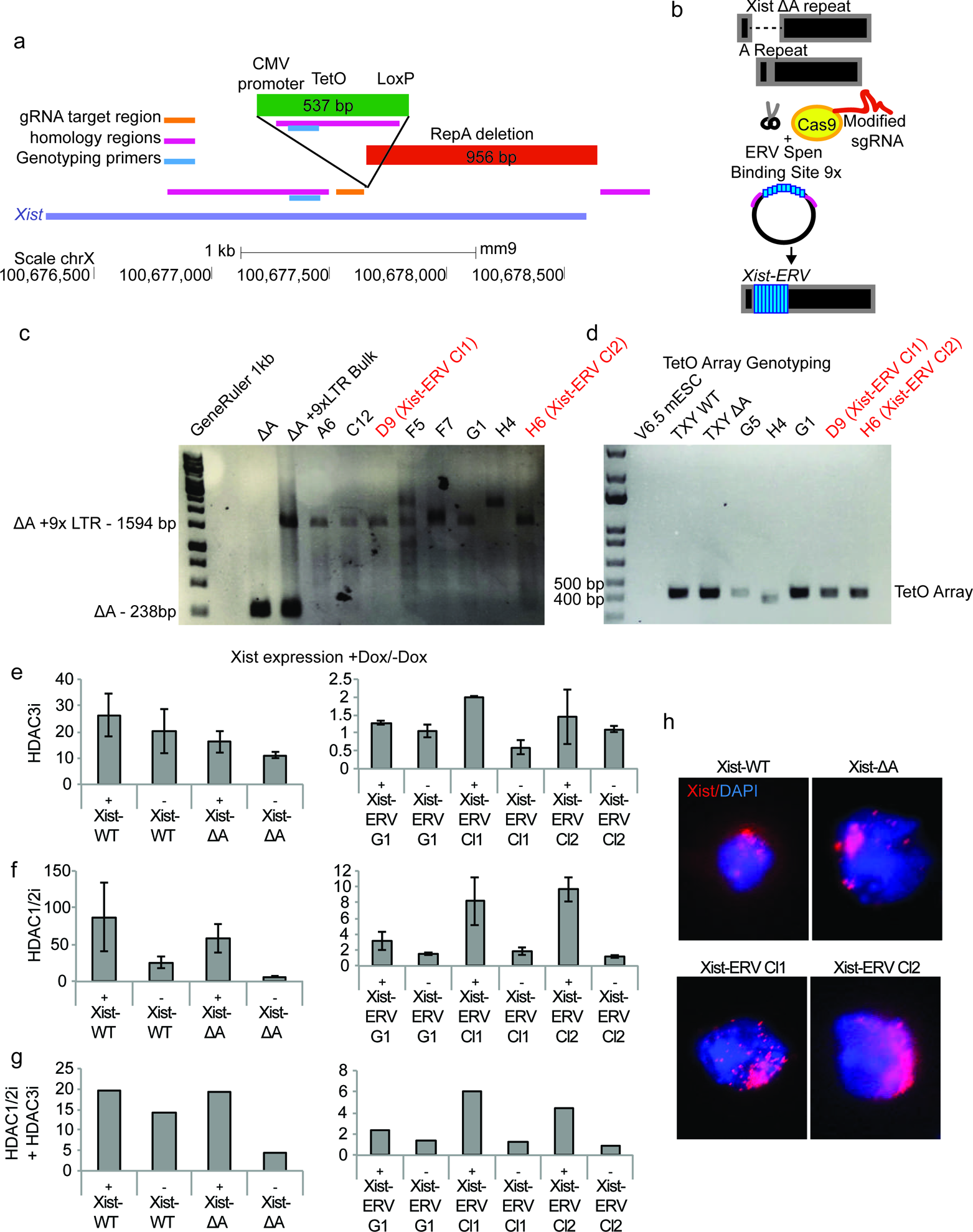
Insertion of an ERV element in the A-Repeat locus. a. Diagram of the *Xist* 5’ end showing the location of the A-Repeat deletion^8^, the CMV promoter and TetO array, and the gRNA target region, homology regions, and genotyping primers for CRISPR-Cas9-directed HDR experiments. b. Diagram of the HDR strategy for inserting a 9x ERV repeat into the A-Repeat locus. Briefly, we used Cas9 to cut at the site where the A-Repeat has been deleted. We then added a donor plasmid containing 2 homology arms to this site flanking a 9x ERV Spen binding site (154 bp per repeat). c. Genotyping showing the insertion of the 9x ETn repeat into the *Xist* locus. Clones shown in red (D9 Xist-ERV Cl1 and H6 Xist-ERV Cl2) are used for further analysis. d. Genotyping results for PCR of the TetO array in 9x ERV knock-in cell lines. The band is at the size of the TetO array knocked into the Xist-WT and Xist-ΔA cell lines. All clones used in this study show a preserved TetO array, indicating that there has not been recombination at this locus. One clone showed a band at a lower height (H4), indicating recombination, but this was not used for further analysis. Primers span from the CMV promoter across the TetO array. e. qPCR data showing *Xist* expression in +Dox relative to –Dox conditions for Xist-WT, Xist-ΔA and Xist-ERV cell lines treated with or without HDAC3i for 24 hours before Doxycycline addition. Data is normalized to *ActB* and then to –Dox. Error bars represent the standard deviation across 4 independent replicates. + and – below the graph refer to whether the cells were treated with HDAC inhibitor. DMSO was used as a control for –HDACi conditions. f. Same as in e for HDAC1/2 inhibitor. g. Same as in e for combined HDAC1/2i and HDAC3i. 2 replicates were analyzed in this experiment. h. FISH for *Xist* RNA in Xist-WT, Xist-ΔA, and Xist-ERV cell lines. Representative nuclei from each cell line are shown. *Xist* FISH signal is in red and DAPI in blue.

**Extended Data Figure 11:**
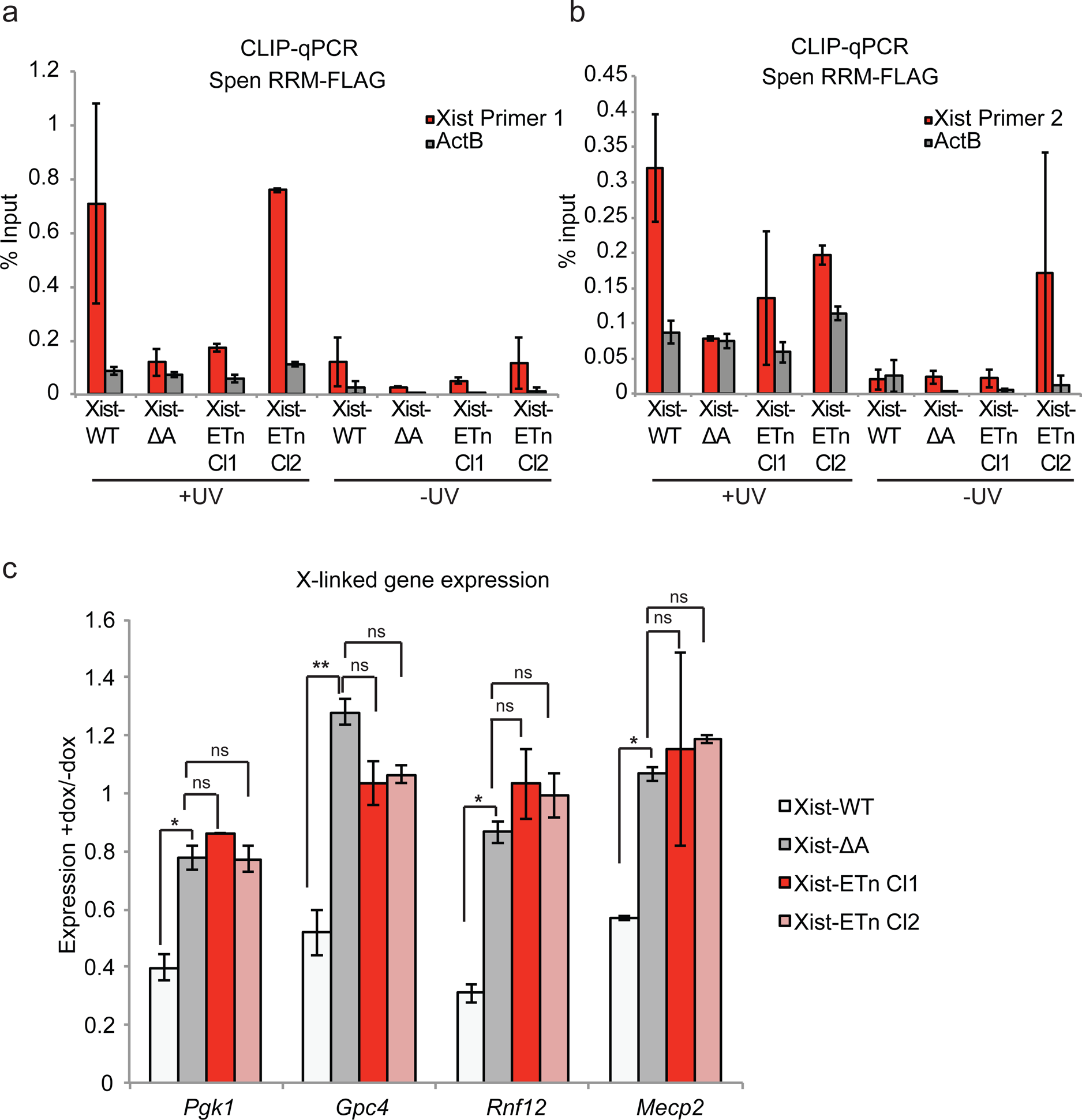
Characterization of *Xist-ERV* function. a. CLIP-qPCR showing recruitment of Spen RRM2-4 by *Xist-ERV* RNA. Each experiment was performed in duplicate. Error bars indicate the standard error. Values for each cell line are normalized to the input sample (as percent input). b. Same as in a for a second set of *Xist* qRT-PCR primers. c. qPCR data showing the silencing of X-linked genes upon *Xist* expression in Xist-WT, Xist-ΔA and Xist-ERV cell lines. All experiments were performed in the presence of HDAC1/2 inhibitor. Expression is normalized first to *ActB* and then to *Xist* level in –dox samples. Error bars show the standard error for 3 replicates.

## Methods

### Cell Lines and Culture

HATX WT and Spen KO clone 2 and clone 3 cell lines were gift of A. Wutz and are described in detail in Monfort et al., 2015^1^. TXY WT (Xist-WT) and TXY deltaA (Xist-ΔA) cell lines were gift of E. Heard. mESCs were cultured in serum/LIF conditions (Knockout DMEM (GIBCO 10829018), 1% Penicillin/streptomycin, 1% GlutaMax, 1%non-essential amino acids, beta-mercaptoethanol, Hyclone fetal bovine serum (Fisher SH3007103), and 0.01% LIF (ESGRO ESG1107). Cells were cultured on 0.2% gelatin-covered plates and passaged every 2-3 days with trypsin as needed.

### *Xist* Induction

For *Xist* induction experiments, mESCs were plated at a density of 75,000 cells/well in a 6-well plate. 24 hours after plating, cells were treated with 2ug/mL doxycycline for 48 hours (unless shorter or longer time is indicated in figure legend). Induction of *Xist* was confirmed using qPCR and *Xist* RNA FISH. qPCR primers used for Xist have the following sequences: Forward 5’GCTGGTTCGTCTATCTTGTGGG3’ and Reverse 5’CAGAGTAGCGAGGACTTGAAGAG3’.

### HDAC inhibition experiments

For HDAC inhibition experiments, cells were treated with either HDAC1/2i (BRD6688, Cayman Chemical Company, 1404562-17-9) or HDAC3i (RGFP966, Sigma SML1652) at 10uM final concentration. HDAC inhibition was initiated 24 hours prior to induction of *Xist* with doxycycline and continued for 24 hours during *Xist* induction (48 hours total). –HDACi samples were treated with DMSO as a control.

### *Xist* RNA FISH

Following *Xist* induction, adherent cells on coverslips were fixed using 4% paraformaldehyde for 10 minutes. Cells were permeablized in 0.5% triton for 10 minutes on ice and then stored in 70% EtOH. After dehydration in EtOH, cells were rehydrated in wash buffer (2xSSC + 10% formamide) for 5 minutes. Probe (Stellaris mXist 570, SMF-3011-1) was then added at a concentration of 125 nM (in wash buffer + 10% Dextran Sulfate) and incubated overnight at 37’C. Following washes in wash buffer + formamide, coverslips were mounted on slides with Vectashield + DAPI.

### NPC Differentiation

NPC differentiation from mESCs was performed as previously described^3^. Briefly, mESCs were plated on gelatin-coated plates in N2B27 medium for 7 days. On day 7, cells were dissociated with Accutase and cultured in suspension in N2B27 medium with FGF and EGF (10 ng/ml, each). On day 10, embryoid bodies were plated onto 0.2% gelatin-coated plates and allowed to grow until 14 days. At 14 days, cells were stained with anti-Nestin (Millipore, MAB353) antibody to confirm NPC identity. For qRT-PCR RNA was extracted from Trizol using the RNEasy Mini kit (Qiagen). RNA was then used directly for qRT-PCR using the following primers. Oct4-F: 5’ TTGGGCTAGAGAAGGATGTGGTT3’, Oct4-R: 5’ GGAAAAGGGACTGAGTAGAGTGTGG3’ ActB-F: 5’TCCTAGCACCATGAAGATCAAGATC3’, ActB-R: 5’CTGCTTGCTGATCCACATCTG3’.

### Immunofluorescence

Immunofluorescence for Oct4 and Nestin was performed on adherent cells fixed with 4% paraformaldehyde. After fixation, cells were permeablized with 0.5% PBS/Triton for 10 minutes on ice. Following permeablization, cells were blocked in 10% FBS in 0.1% PBS/Tween for 1 hour at room temperature. After blocking, cells were incubated for 2 hours in primary antibody (Nestin: Millipore, MAB353, Oct4: Santa Cruz sc-8629), washed in 0.1% PBS/Tween and then incubated for 2 hours in secondary antibody (Goat anti-mouse 488 Life Technologies ab150113, Rabbit anti-goat 488 Life Technologies A27012). After washing, cells were stained with DAPI and imaged at 40x magnification.

### Chromatin Immunoprecipitation

Cells were fixed with 1% formaldehyde for 10 minutes at room temperature and subsequently quenched with 0.125M glycine. Cells were then snap frozen and stored at −80’C. Cells were then lysed (50mM HEPES-KOH, 140 mM NaCl, 1mM EDTA, 10% glycerol, 0.5% NP-40, 0.25% Triton X-100) for 10 minutes at 4’C. Nuclei were lysed (100 mM Tris pH 8.0, 200mM NaCl, 1mM EDTA, 0.5 mM EGTA) for 10 minutes at room temperature. Chromatin was resuspended in sonication buffer (10 mM Tris pH 8.0, 1mM EDTA, 0.1% SDS) and sonicated using a Covaris Ultrasonicator to an average length of 220 bp. For all ChIPs, 5 million cells per replicate were incubated with 5 ug (Histone) or 7.5 ug (Oct4) antibody overnight at 4’C (antibodies: H3K9me3: abcam AB8898; H3K27Ac: Active Motif 39133; H3K4me3: Active Motif 39159; Oct4: Santa Cruz sc-8629). Antibody-bound chromatin was incubated with Protein G Dynabeads (Invitrogen, 10004D) for 4 hours at 4’C and eluted in Tris buffer (10mM Tris pH 8.0, 10mM EDTA, 1% SDS). Crosslinks were reversed by incubation overnight at 65’C followed by treatment with 0.2 mg/mL proteinase K (Life Technologies, AM2548) and 0.2 mg/mL RNAse A (Qiagen). DNA was purified using Qiagen Minelute Columns (Qiagen, 28006). For library preparation, 4 ng ChIP DNA was incubated with transposase (Illumina, Fc-121-1030) for 10’ at 55’C. DNA was then amplified using Nextera barcoding adapters and sequenced on an Illumina Nextseq (2×75 bp reads).

### ChIP-seq analysis

ChIP-seq libraries for histone modifications were sequenced on an Illumina NextSeq with paired-end 75 bp reads. Reads were mapped to the mm9 genome build using Bowtie2^4^. Unique and coordinated mapped paired reads were kept and duplicate reads were then removed using Picard Tools.

Peaks were called using MACS2 with a q-value cutoff of 0.01 with input as background^5^. Peaks called from two technical replicates were compared by IDR package and further filtered by the Irreproducible Discovery Rate(<0.05)^6^. Reproducible peaks from WT and KO conditions were merged and reads in these peaks were counted across all libraries. Differential peaks were called using reads counts from reproducible peaks with DESeq2(fold change >=2 and Padjust <0.01)^7^. Bigwig files were generated from deduplicated bam files using bedtools for visualization^8^.

### ATAC-seq

ATAC-seq library preparation was performed exactly as described in Buenrostro et al., 2013^9^. Briefly, ESCs were dissociated using Accutase (SCR005,Millipore). 50,000 cells per replicate (2 replicates per clone) were incubated with 0.1% NP-40 to isolate nuclei. Nuclei were then transposed for 30’ at 37’C with adapter-loaded Nextera Tn5 (Illumina, Fc-121-1030). Transposed fragments were directly PCR amplified and sequenced on an Illumina NextSeq 500 to generate 2x 75bp paired-end reads.

### ATAC-seq analysis

Reads were trimmed using CutAdapt^10^ and mapped to the mm9 genome build using Bowtie2^4^. Unique and coordinated mapped paired reads were kept and duplicate reads were then removed using Picard Tools. Peaks were called using MACS2 with a q-value cutoff of 0.01 and no shifting model^5^. Peaks called from two technical replicates were compared by IDR package and further filtered by the Irreproducible Discovery Rate(<0.05)^6^. Reproducible peaks from WT and KO conditions were merged and reads in these peaks were counted across all libraries. Differential peaks were called using reads counts from reproducible peaks with DESeq2(fold change >=2 and P-adjusted <0.01)^7^. To calculate *P* values in DESEq2, for each gene the counts were modelled using a generalized linear model (GLM) of negative binomial distribution among samples. The two-sided Wald statistics test was processed for significance of the GLM coefficients. The Benjamini-Hochberg correction was applied to all p-values to account for multiple tests performed. This gives the final adjusted p-values used for assessing significance.

### ATAC-seq analysis for X chromosome silencing

ATAC-seq reads were trimmed and mapped as described above. Peaks were called in all samples (+ and – doxycycline) and bedtools multicov was used to count the number of reads falling within each peak in each sample. In order to avoid bias in normalization in the case of a global loss of accessibility on the X chromosome, we first normalized count data using all peaks genome-wide. We then calculated the ratio of average normalized reads in +dox/-dox samples at each peak on the X chromosome.

### RNA-seq

RNA-seq library preparation was performed using the TruSeq Stranded mRNA Library Prep Kit (Illumina, RS-122-2102). In place of poly-A selection, we performed Ribosomal RNA depletion using the RiboMinus Eukaryote System v2 (Life Tech. A15026). Each library was prepared from 200ng starting RNA, and 2 replicates were made for each cell line. RNA-seq libraries were sequenced on an Illumina HiSeq 4000 with paired 2×75 bp reads.

### RNA-seq analysis

RNA-seq reads were mapped to the mm9 genome using STAR^11^. Following read alignment, reads were assigned to transcripts using FeatureCounts in R (RSubread)^12^. GTF file from gencode was used for feature assignment (https://www.gencodegenes.org/mouse_releases/1.html). Differential genes were called using DESeq2 (fold change >=2, P-adjusted<0.001)^7^. For GO term analysis of RNA-seq data, GOrilla was used^13^.

### Enrichment of genomic loci in ATAC-seq and ChIP-seq data

HOMER was used for enrichment of genomic locations in ATAC-seq and ChIP-seq data^14^. Specifically, we used the annotatePeaks.pl script with the –genomeOntology option.

### *In vivo* irCLIP-seq

For in vivo irCLIP-seq experiments, V6.5 male mESCs, or *Spen* KO Clone 2 mESCs were transduced with a lentivirus carrying Spen RRMs 2-4 with the SV40 nuclear localization signal, 2x HA tags, and 3x FLAG tag (“SpenRRM-FLAG”; pLVX-EF1a-SV40NLS-SpenRRM234-2xHA-3xFLAG-IRES-zsGreen). Following transduction, GFP+ cells were FACS sorted and single colonies were picked. Western blotting was used to confirm the expression of SpenRRM-FLAG at 37 kDa. irCLIP was performed exactly as described^15^. Briefly, 2 million mESCs were UV-crosslinked (254nM UV-C) at 0.3J/cm^2^, and then lysed (1% SDS, 50mM Tris pH7.5, 500 mM NaCl) and sonicated using a Bioruptor (Diagenode) (six cycles 30” on 45” off). Clarified lysates were incubated overnight with Protein A dynabeads conjugated to mouse anti-FLAG (Sigma Aldrich F3165) or anti-IgG antibody overnight at 4C. Following IP, beads were washed in high stringency buffer (15mM Tris-HCl, pH 7.5; 5mM EDTA; 1% Triton X-100; 1% Na-deoxycholate; 0.001% SDS; 120mM NaCl; 25mM KCl), high salt buffer (15mM Tris-HCl, pH 7.5; 5mM EDTA; 1mM Triton X-100; 1% Na-deoxycholate; 0.001% SDS; 1000mM NaCl), and low salt buffer (15mM Trish-HCl, pH 7.5; 5mM EDTA). After washing, RNA was digested using RNase I at 1:2000. RNA ends were then dephosphorylated and IR-conjugated linker (/5Phos/AGATCGGAAGAGCGGTTCAGAAAAAAAAAAAA/iAzideN/AAAAAAAAAAAA/3Bio/) was ligated overnight. Following ligation, samples were run on a 4-12% Bis-Tris gel, transferred to nitrocellulose and imaged using an Odyssey LiCOR scanner. RNA-protein complexes were eluted from nitrocellulose and treated with proteinase K to eliminate protein. Following RNA elution, Oligo(dT) Dynabeads were added to capture the polyadenylated irCLIP adapter. Next, the TruAmp sequencing adapter was added and RNA was reverse transcribed using SuperScript IV. Following reverse transcription, DNA was captured with streptavidin beads, circularized on the bead, and then PCR amplified. irCLIP libraries were then purified using PAGE gel purification and sequenced on an Illumina NextSeq using single-end 75 bp reads.

### Expression and purification of wild-type and mutant Spen RRMs 2-4 for *in vitro* irCLIP RNA binding

The following constructs for SPEN (Split Ends protein), SHARP (human homolog, SMRT/HDAC1 Associated Repressor Protein), were designed and cloned into pET28a for E.coli expression. SPEN-1= 6His-GST-TEV-RRM 2,3,4-SG-Flag SPEN-4= 6His-GST-TEV-RRM 2,3,4-RRM3mutant-SG-Flag DNA for each was transformed into BL21 (DE3) Star competent cells, the transformation mix plated on LB-Kan agar and single colonies obtained the following day. 4 x 50ml overnight pre-cultures were setup with single colonies for each construct in Terrific Broth (Teknova) + Kanamycin (50ug/ml) (TB-Kan), at 37C, 250rpm. Pre-cultures were used to inoculate 1li volumes of TB-Kan expression media at 10ml/li (gives a starting OD600=0.15). Expression cultures were incubated at 37C, shaken at 250rpm, for 4hrs then chilled to 18C for 1hr and induced overnight with 0.25mM IPTG at 18C. Cultures were harvested and resulting cell pellets were stored frozen at −80C. Frozen cell pellets were resuspended in 5ml/g of Lysis Buffer(50mM HEPES;pH 7.5, 300mM NaCl, 20mM Imidazole, 0.1% Triton X-100, DNase I (1ug/ml), Lysozyme (1ug/ml), 5mM 2-ME, 2 x Roche protease Inhibitor tablet/50ml and 1mM PMSF). Resuspended pellets (in suspensions) were homogenized by Polytron and lysed by 2 rounds through a Microfluidizer (18000psi). Lysates were clarified by centrifugation at 33000 x g for 60mins, the supernatant extracted and subjected to batch binding with 5ml of NiNTA (Qiagen) for 2hrs, 4C with end-to-end rotation. Resin was washed with 20Cv’s of Lysis Buffer containing 20mM Imidazole, followed by elution of bound protein with 3 x 1CV of Elution Buffer (Lysis Buffer + 250mM Imidazole). Eluted fractions were pooled and dialysed overnight against 4li of 50mM Hepes;pH 7.5, 300mM NaCl, 1mM TCEP with the addition of TEV protease to effect in-situ cleavage of the N-terminal 6His-GST tag. TEV cleaved solution was passed through a 0.5ml NiNTA and the flow-through collected and concentrated 3-5 fold to give 4-10mg/ml samples which were aliquotted and stored frozen at −80C.

### *In vitro* irCLIP RNA binding

*In vitro* irCLIP-seq was performed similarly to *in vivo* irCLIP with the following modifications. First, nuclear RNA was isolated as previously described^16^. Briefly, cells were pelleted and resuspended in Hypotonic Lysis Buffer (10 mM Tris pH 7.5, 10mM NaCl, 3mM MgCl_2_, 0.3% NP-40 and 10% glycerol) with 100 Units SUPERase-In. Cells were incubated on ice for 10 minutes, vortexes, and centrifuged for 3 minutes at 1,000xg at 4C. The pelleted nuclei were then washed 3x in hypotonic lysis buffer and centrifuged for 2 minutes at 200xg at 4C. Pellet was then resuspended in TRIzol and RNA extraction was carried out per the manufacturer’s instructions. 20 μg nuclear RNA was then fragmented (Ambion, AM8740), and incubated with 5 μg recombinant Spen RRMproteins (RRM2-4 WT or RRM2-4:RRM3 Mt) at 4C for 2 hours. RNA-protein complexes were then transferred to a 3cm cell culture dish on ice and crosslinked (254nM UV-C) at 0.3J/cm^2^. Following crosslinking, RNA-protein complexes were incubated with antibody-bound Protein A Dynabeads (anti-FLAG [Sigma Aldrich F3165] or anti-IgG) at 4C for 2 hours. Following IP, beads were washed with IPP buffer (50 mM Tris-HCl pH 7.4, 100 mM NaCl, 0.05% NP-40, 5mM EDTA), low salt buffer (10 mM Tris-HCl pH7.4, 50 mM NaCl, 1mM EDTA, 0.1% NP-40), and high salt buffer (10 mM Tris-HCl pH7.4, 500 mM NaCl, 1 mM EDTA, 0.1% NP-40, 0.1%SDS). After washes, the protocol was performed exactly as in *in vivo* irCLIP from dephosphorylation to membrane imaging(above).

### Analysis of expressed repeats in RNA-seq data

To identify expressed transposable elements in RNA-seq data, we counted all unique reads mapping to the database of repeat element loci from RepeatMasker for the mm9 genome. Bedtools multicov was used to count the number of reads per sample in each element. Read counts were then converted to fpkm. All elements with an FPKM >=1 in all samples were considered expressed and used for downstream analysis.

### ERV knock-in at the *Xist* A repeat

For Xist rescue experiments with ETn elements, we utilized the TXY ΔA cells which harbor a doxycycline inducible Xist transgene with the A Repeat region deleted. We designed donor plasmids for homology directed repair containing homology arms to the transgenic Xist present in TXY ΔA cells that contain a 154 bp portion of an ETn that binds to Spen, repeated 9x. Homology arms flanking the ETn insertions were cloned into the PCR4-Blunt TOPO vector (Thermo Fisher K287520). Cells were then nucleofected with a Cas9 RNP complex directed to the A Repeat region. 10 ug Cas9 protein (PNA Bio CP01-20) was mixed with 10 ug modified guideRNA (Synthego, GCGGGATTCGCCTTGATTTG) and 20 ug HDR donor vector. Cells were nucleofected with RNP complex and donor vector using the Amaxa Nucleofector (Lonza VPH-1001). Clonal colonies were picked and genotyped by PCR followed by Sanger sequencing. PCR of the Tet operator array was done using the following primers: TetO-F: 5’CCTACCTCGACCCGGGTACC3’, TetO-R: 5’GGCCACTCCTCTTCTGGTCT3’.

### RIP-qPCR

TXY WT, TXY ΔA, and TXY-ERV Clones 1 and 2 were transduced with a lentivirus carrying Spen RRMs 2-4 with the SV40 nuclear localization signal, 2x HA tags, and 3x FLAG tag (“SpenRRM-FLAG”; pLVX-EF1a-SV40NLS-SpenRRM234-2xHA-3xFLAG-IRES-zsGreen). Following transduction, GFP+ cells were FACS sorted and expression of the RRM-FLAG construct was confirmed by Western Blot using an anti-FLAG antibody (Sigma F3165). SpenRRM-FLAG expressing cells were treated with 10 mM HDAC1/2 inhibitor (Cayman Chemical Company, BRD6688, 1404562-17-9) for 24 hours, followed by treatment with HDAC1/2i + 2 mg/mL doxycycline for 24 hours. 5 million cells were then washed in PBS and lysed in RIP lysis buffer (50 mM Tris pH 8.0, 100 mM NaCl, 5 mM EDTA, 0.5% NP-40, 1 cOmplete Protease inhibitor tablet) in the presence of RNAse inhibitor (Thermo Scientific, RiboLock RNAse Inhibitor, EO0382). Cells were sonicated on a Covaris (Fill level 10, Duty Cycle 5%, PIP 140 Watts, 200 cycles/burst, 5×1minute/sample), and debris was pelleted for 15 minutes at 21,000xg at 4’C. Cleared supernatant was then added to 100 uL FLAG magnetic beads (Sigma, anti-FLAG M2 Magnetic Beads, M8823). After 2 hours incubation at room temperature, beads were washed 3 times in cell lysis buffer and resuspended in Trizol. RNA was extracted using the RNEasy mini kit (Qiagen, 74106). qRT-PCR was run on a Roche Lightcycler 480 using the Brilliant II SYBR Green qPCR Master Mix (Stratagene, 600804) using the following primers: Xist-F: 5’GCTGGTTCGTCTATCTTGTGGG3’, Xist-R: 5’CAGAGTAGCGAGGACTTGAAGAG3’, ActB-F: 5’TCCTAGCACCATGAAGATCAAGATC3’, ActB-R: 5’CTGCTTGCTGATCCACATCTG3’.

### Multimapping for repeat-derived reads in ChIP-seq and ATAC-seq data

After finding that SPEN regulates repeat regions, we re-analyzed the ChIP-seq and ATAC-seq data to include repetitive regions, which are usually removed in the standard analysis. We allowed each multimapping read to randomly assign to one place in the genome, without filter reads by mapping quality. The differential peaks is consistent to the results from unique mapped reads. We used the alignment files with multiple mapped reads for all the meta-analysis to avoid losing information.

### Evolutionary analysis of Spen

Multiple alignments of the SPEN CDS sequence were download from UCSC, including human, chimpanzee, rhesus, mouse, dog, opossum, platypus, chicken, zebrafish. We separated the CDS of SPEN into three domain regions (RID, SPOC and RRM).

We estimated the number of nonsynonymous substitutions per nonsynonymous site (*dN*), the number of synonymous substitutions per synony-mous site (*dS*), and the *dN*/*dS* ratio, for each region using maximum likelihood as implemented in the codeml program of the PAML software package (Yang 2007)^18^. Model 0 were used to estimate the global evolution rate and model 1 were used to estimate branch specific evolution rate.

### Integrated analysis of ChIP-seq, RNA-seq, and ATAC-seq data

Differentially expressed genes were identified from RNA-seq data. The 1kb region around the promoter of differentially expressed genes were used to count the signal from ChIP-seq and ATAC-seq data. The fold change of gene expression, histone modification and chromatin accessibility between WT and *Spen* KO were compared using a R package, named tableplot^20^. The average plots and heatmap from histone modification, RNA expression, accessibility for interested regions were made using R package ngsplot^21^.

### Repeat Divergence analysis

The divergence scores were download from repeatMasker through UCSC genome browser. We separated the ETn family by repeats with binding sites detected from irCLIP or not and the divergence score from mismatches (as a percentage) were compared.

### icSHAPE data analysis

The icSHAPE data from Sun et al. were downloaded from GEO under accession GSE117840. The sequence data reads were collapsed to remove PCR duplicates and trimmed to remove 3’ adapters following the icSHAPE computational pipeline^22^. The processed reads were mapped to the expressed repeat sequence using bowtie2 with parameters suggested by Flynn et al^22^. The icSHAPE score were then calculated following the previous description. Repeat sequence with read depth more than 75 were kept for the following analysis. The icSHAPE score of SPEN binding sites and their flanking region(+/-20bp) were extracted, The aggregated icSHAPE activity score were plotted for each base across non-overlapped binding locations on the Ent repeats.

### irCLIP sequencing data processing

Sequencing output reads were processed by bbmap to remove the duplication on fastq level. Remained reads were trimmed off the 3’ solexa adapter and against the sequencing quality q20 by cutadapt (version 2.4). Trimmed reads were mapped first to the RNA biotypes with high repetitiveness by bowtie2 (version 2.2.9) to our custom built indexes: rRNAs (rRNAs downloaded from Ensembl NCBI37/mm9 and a full, non-repeat masked mouse rDNA repeat from GenBank accession No.BK000964), snRNAs (from Ensembl NCBI37/mm9), miscRNAs (from Ensembl NCBI37/mm9), tRNAs (from UCSC table browser NCBI37/mm9), retroposed genes (RetroGenes V6) and RepeatMasker (from UCSC table browser NCBI37/mm9). Remained reads were mapped to the mouse genome NCBI37/mm9 by STAR (version 2.7.1a) with junction file generated from mRNAs and lncRNAs of the Genocode NCBI37/mm9 GTF file. Only reads uniquely mapped to the mouse genome were included in the down-stream analysis. Mapping output files were transformed into bam files (samtools/1.9) and bed files (bedtools/2.27.1) for downstream analysis.

### SPEN binding cluster identification

The RBP binding loci as suggested by the irCLIP method, was defined as one nucleotide shift to the 5’ end of each mapped read. Each loci was extended five nucleotides both up and downstream into a 11-nt-long local interval. Only intervals overlapped between two biological replicates were considered reliably detected and were included in the downstream analysis. Overlapping intervals were merged into one cluster. Five nucleotides were trimmed from each side of the cluster to shape the final cluster boundary. Cluster length is required to be longer than one nucleotide for downstream analysis.

SPEN binding clusters on mouse genome and on repeat-regions were identified respectively. Cluster annotations were processed against the Genocode NCBI37/mm9 GTF file and RepeatMasker.

For each experiment condition of each cell type, only clusters with quantification tenfold higher than the observation in its corresponding IgG samples, would be included for the downstream analysis.

### Gene/repeat element level quantification of SPEN binding

Quantification of SPEN binding on each expressed gene or expressed repeat element was calculated as the total amount of read in all binding clusters covering the gene/repeat element. The quantifications were then normalized by gene/repeat expression level.

### SPEN binding RNA motif analysis

RNA motif enrichment analysis were process on SPEN binding clusters on mouse genome and on repeat-regions respectively (Homer version 4.10). Strand-specific sequences defined by SPEN irCLIP clusters were searched for short sequences elements 5-mer, 6-mer, 7-mer and 8-mer in each analysis of hypergeometric enrichment calculations. The searching background was set as expressed transcriptome and expressed repeat regions.

### RNA 2D-structure

2D-structures were predicted by the RNAstructure tool (version 6.0.1) with the maximum free energy model and according the icSHAPE results. 2D-structures of MMETn repeat1125276, ETnERV repeat355428 and RLTR13G_2D were process by algorithm AllSub. 2D-structure of Xist repeat A region #4 and #5 were processed with algorithm DuplexFold.

### Cloning of wild-type and mutant Spen RRMs 2-4 for *in vitro* biochemistry

Wild type mouse Spen cDNA was kindly provided in a D-TOPO vector from the Guttman lab (Caltech). Mutant Spen cDNA corresponding to RRMs 2-4 (F442A, K471A, Y479A, F481A, K513A) was ordered as a gBlock from IDT (Coralville, IA). Primers containing SalI and XhoI restriction enzyme cut sites on the 5’-end and 3’-end respectively of the cDNA corresponding to RRMs 2-4 (aa 336-644) were used to PCR amplify both wild type and RRM3 Mt cDNA inserts for ligation into a pET28b vector containing an N-terminal 10xHis-SUMO tag using the Quick Ligation kit (NEB, Ipswich, MA). Plasmids containing the cDNA of interest were verified by Sanger sequencing prior to expression (Quintara Biosciences, San Francisco, CA).

### Expression and purification of wild-type and mutant Spen RRMs 2-4 for *in vitro* biochemistry

BL21(DE3) expression cells were transformed with pET28b vector containing the cDNA of the protein of interest. 1 L cultures of 2XYT media were grown at 37°C for 3 hours to an OD_600_ between 0.6-1.2. Cells were flash-cooled on ice for 10min, then 1 mM IPTG was added to induce protein expression overnight at 20°C. Cells were harvested, and pellets were frozen and stored at −20°C until purification. All proteins were subject to a 3-column purification protocol; the wild-type buffers were pH 7.5, while the mutant buffers were pH 8.3 due to differences in predicted pI and anticipated solubility requirements. Specifically, one cOmplete EDTA-free EASYpack protease inhibitor tablet (Roche, Basel, Switzerland) was added to each cell pellet corresponding to 1 L of growth. The tablet and frozen cell pellet were resuspended in 80 mL of lysis buffer (750 mM KCl, 750 mM NaCl, 1 M urea, 50 mM Tris, 10% glycerol, 10 mM imidazole, 1% triton X-100). The cell slurry was sonicated on ice with a Misonix Sonicator 3000 (Misonix, Farmingdale, NY) for 4-6 total minutes in pulses of 15sec followed by 35sec of rest. Membranes and other cellular debris were sedimented away from soluble protein on a Beckman J2-21 large floor centrifuge (Beckman Coulter, Brea, CA). Supernatant containing the recombinant protein was incubated with 10 mL of precleared Ni-NTA beads for 10min at 4°C. Unbound cellular proteins were cleared by gravity flow. Beads were washed twice; once with 150 mL of lysis buffer, and once with 100 mL of modified lysis buffer containing 20 mM imidazole. Protein was then eluted with 50 mL of similar buffer containing 300 mM imidazole. For the washes and elution, beads were resuspended and incubated with the buffer for 10min prior to gravity flow through. The elution was dialyzed in 4 L of size exclusion column buffer (500 mM urea, 270 mM KCl, 30 mM NaCl, 50 mM Tris, 7% glycerol, 1-2 mM DTT) for about 2hrs at 4°C. 1-2 mL of 1 mg/mL in-house His-ULP1 [Mossessova and Lima, Mol. Cell (2000), 865-876] was then added, and SUMO tag cleavage of recombinant Spen protein and further dialysis continued overnight at 4°C. Protein precipitate, when present, was pelleted or filtered prior to flowing over the second Ni-NTA column (11 mL beads precleared in column buffer). The Spen fragment flowed through the second Ni column while the His tagged protease and SUMO expression tag were retained on the resin. Flowthrough containing the cleaved Spen construct was concentrated to 2 mL for injection onto the Superdex 200 size exclusion column. Size exclusion fractions are checked for the presence of purified protein by SDS-PAGE. Fractions containing purified protein are combined, concentrated to between 100 µM −1 mM, aliquoted, flash frozen in liquid nitrogen, and stored at −70°C until binding assays are performed.

### RNA generation

cDNAs of RNAs of interest with a 5’ T7 promoter were ordered as gBlocks from IDT (Coralville, IA). Primers flanking the cDNA containing the EcoRI and BamHI cut sites on the 5’ and 3’ ends respectively were used to PCR amplify the cDNA for restriction enzyme cloning into a pUC19 vector for cDNA amplification. Ligated plasmids were verified using Sanger sequencing (Quintara Biosciences, San Francisco, CA) and utilized for transcription template generation. Standard PCR using a 3’ transcription primer was used to generate *in vitro* transcription template for run-off transcription. For *in vitro* transcription, 300 nM PCR template was incubated at 37°C for 2-4hrs with 4 mM each rNTP in 24 mM Mg^2+^, 40 mM Tris pH 8.0, 10 mM DTT, 2 mM spermidine, 0.01% Triton X-100, IPPase (>0.01U/µL), and in-house T7 polymerase (Batey Lab, CU Boulder). Transcription reactions were gel purified on a 5-12% acrylamide 8M urea 1X TBE slab gel. Reaction products were UV shadowed, appropriate sized bands were excised, and RNA was extracted through a 0.5X TE crush and soak (4°C, 1hr). Gel bits were filtered and RNA was buffer exchanged in 0.5X TE until the calculated residual urea was less than 10 µM. RNA was concentrated prior to use; purity and concentration were assessed by the A_260_ and the A_260_/A_280_ ratio. Concentrations were calculated using the extinction coefficient provided for each RNA sequence by the Scripps extinction coefficient calculator (https://www.scripps.edu/researchservices/old/corefac/biopolymercalc2.html).

### FTSC 3’-end labeling of RNA

350 pmol of purified RNA were treated with 20 mM NaIO_4_ for 20min in the dark at room temperature. Excess NaIO_4_ was precipitated using 26 mM KCl (10min on ice followed by a hard spin). Supernatant containing the RNA was then ethanol precipitated with the aid of 1 µg glycogen. The RNA pellet was washed 1x with 70% ethanol, and then resuspended in 100 µL of freshly made labeling solution (1.5 mM FTSC, 10% DMF, 100 mM NaOAc pH 5.2) and reacted at 37°C in the dark for 1-2hrs. 280 µL of cold 100% ethanol was then added to precipitate the RNA (ice for ≥ 30min). The labeled RNA pellet was then washed thoroughly at least 4X with 400 µL of 70% ethanol, resuspended in 30 µL of water, and run through a Sephadex MicroSpin G-25 column (GE Healthcare) to remove excess FTSC. Labeled products were qualitatively assessed for purity and labeling efficiency via denaturing gel analysis; the Cy2 filter on the Typhoon imager (GE Healthcare) was used to visualize the attached fluorophore. A_260_ measurements post-labeling were used to determine RNA concentration. Labeling efficiencies typically allowed for the use of RNA in binding assays between 1-3 nM. FTSC-labeled RNA was stored in dark amber tubes at −20°C until use.

### Fluorescence anisotropy (FA) binding assays

Wild-type and mutant Spen binding were performed at pH 7.5 and 8.3 respectively. RNA concentrations were held as low as possible while still obtaining enough fluorescent signal (1-3 nM). Briefly, purified labeled RNA was snap cooled in 1X binding buffer (45 mM KCl, 5 mM NaCl, 50 mM Tris, 5% glycerol) at 2X the final concentration, and likewise protein titrations were performed separately in 1X binding buffer at 2X final concentration. Protein and RNA were then mixed in a 1:1 volume ratio in a 20 µL reaction and were allowed to come to equilibrium at room temperature in the dark for 40-60min. Flat bottom low flange 384-well black polystyrene plates (Corning, Corning, NY) were used. Perpendicular and parallel fluorescence intensities (I_⊥,_I_ǁ‖_) were measured using a ClarioStar Plus FP plate reader (BMG Labtech). Anisotropy values were calculated for each protein titration point where anisotropy = (I_ǁ‖_ -I_⊥_)/(I_ǁ‖_ + 2*I_⊥_). Protein concentration was plotted versus associated anisotropy, and data were fit to the simplified binding isotherm (anisotropy ∼ fraction bound = S*(P/(K_D_+P))+O with KaleidaGraph where S and O were saturation and offset respectively, and P was the protein concentration.

### ^32^P 5’-end labeling of RNA

50 pmol of RNA were treated with calf intestine phosphatase (NEB) for 1hr at 37°C. RNA was purified via phenol chloroform extraction followed by ethanol precipitation. RNA was resuspended and γ-ATP was added to the 5’ end of the RNA with T4 polynucleotide kinase (NEB) for 30min at 37°C. Water was added to dilute the RNA to 1 µM, and the reaction was then run through a Sephadex MicroSpin G-25 column (GE Healthcare) to remove excess γ-ATP. ^32^P-labeled RNA was stored at −20°C.

### Competition assay

Wild-type Spen and ^32^P-labeled A repeat concentrations were kept constant at 10 µM and 1 µM respectively, and the competition was done at 50 mM Tris (pH 7.5), 45 mM KCl, 5 mM NaCl, and 2.5% glycerol. Different amounts of unlabeled competitor RNA were added, and the amounts were relative to the amount of labeled RNA. Briefly, RNAs were snap cooled in RNA buffer (50 mM Tris (pH 7.5), 45 mM KCl, 5 mM NaCl). Labeled RNA was snap cooled at 4X the final concentration, while competitor RNA was snap cooled at 2X the final concentration. Protein was diluted in protein buffer (50 mM Tris (pH 7.5), 45 mM KCl, 5 mM NaCl, 10% glycerol) to 4X the final concentration. 2.5 µL of 4X protein, 2.5 µL of 4X labeled RNA, and 5 µL of 2X competitor RNA were allowed to come to equilibrium at room temperature for 40-60min. The reactions were then loaded onto a pre-run 5% acrylamide 0.25X TBE mini native gel at 200V. The gel was run at room temperature at 200V for 10-15min, dried, then exposed overnight to a phosphoscreen. Phosphoscreen images were taken on the Typhoon (GE Healthcare) at 50 µm resolution.

### List of RNA sequences used in direct binding and competition

A-repeat from Xist (85 nt): GGAACTCGTTCCTTCTCCTTAGCCCATCGGGGCCATGGATACCTGCTTTTAACCTTTCTCGGTCCATCGGGACCTCGGATACCTG
SRA RNA (82 nt): GGAGCAGGTATGTGATGACATCAGCCGACGCCTGGCACTGCTGCAGGAACAGTGGGCTGGAGGAAAGTTGTCAATACCTGTA
ID 205 (86 nt): GAUUCCAGGCAGAACUGUUGAGCAUAGAUAAUUUUCCCCCCUCAGGCCAGCUUUUUCUUUUUUUAAAUUUUGUUAAUAAAAGGGAG
ID 509 (114 nt): GGUUAAAACUGCUAUUGUUCCAUUGACUGCAGCUUGCAGUUUGAUUUCAAAUUUAAGAUCUUUAAUUCACCUGUAUACUGUAAUUAAGAUAAUUACAAGAGUAAUCAUCUUAUG
ID 495 (113 nt): GAAAUUUCUCUCUGGGCCUUAUAGCAGGAGUACUCUGUUCCCUUUUGUGUCUUGUCUAAUGUCCGGUGCACCAAUCUGUUCUCGUGUUCAAUUCAUGUAUGUUCGUGUCCAGU
ID 411 (109 nt): GGUACCUUAAAUCCUUGCUCUCACCCAAAAGAUUCAGAGACAAUAUCCUUUUAUUACUUAGGGUUUUAGUUUACUACAAAAGUUUCUACAAAAAAUAAAGCUUUUAUAA
ID 302 (111 nt): GAUCAGAGUAACUGUCUUGGCUACAUUCUUUUCUCUCGCCACCUAGCCCCUCUUC UCUUCCAGGUUUCCAAAAUGCCUUUCCAGGCUAGAACCCAGGUUGUGGUCUGCUGG

